# CRY1-CBS binding regulates circadian clock function and metabolism

**DOI:** 10.1101/2020.01.09.898866

**Authors:** Sibel Cal-Kayitmazbatir, Eylem Kulkoyluoglu-Cotul, Jacqueline Growe, Christopher P. Selby, Seth D. Rhoades, Dania Malik, Hasimcan Oner, Hande Asimgil, Lauren J. Francey, Aziz Sancar, Warren D. Kruger, John B. Hogenesch, Aalim Weljie, Ron C. Anafi, Ibrahim Halil Kavakli

**Author notes:** Correspondence to. &.

## Abstract

Circadian disruption influences metabolic health. Metabolism modulates circadian function. However, the mechanisms coupling circadian rhythms and metabolism remain poorly understood. Here we report that Cystathionine β-synthase (CBS), a central enzyme in one-carbon metabolism, functionally interacts with the core circadian protein Cryptochrome1 (CRY1). In cells, CBS augments CRY1 mediated repression of the CLOCK/BMAL1 complex and shortens circadian period. Notably, we find that mutant CBS-I278T protein, the most common cause of homocystinuria, does not bind CRY1 or regulate its repressor activity. Transgenic *Cbs*^Zn/Zn^ mice, while maintaining circadian locomotor activity period, exhibit reduced circadian power and increased expression of E-BOX outputs. CBS function is reciprocally influenced by CRY1 binding. CRY1 modulates enzymatic activity of the CBS. Liver extracts from *Cry*1^−/−^ mice show reduced CBS activity that normalizes after the addition of exogenous wild type (WT) CRY1. Metabolomic analysis of WT, *Cbs*^Zn/Zn^, *Cry*1^−/−^, and *Cry2*^−/−^ samples highlights the metabolic importance of endogenous CRY1. We observed temporal variation in one-carbon and transsulfuration pathways attributable to CRY1 induced CBS activation. CBS-CRY1 binding provides a post-translational switch to modulate cellular circadian physiology and metabolic control.

## Introduction

The circadian clock modulates numerous behaviors and physiologic functions through periodic transcriptional regulation [1]. Alertness, memory, heart rate, blood pressure, and immune responses are all clock-controlled [2–4]. Additionally, genetic and epidemiologic studies have linked clock disruption with various adverse metabolic phenotypes [5, 6]. Dietary challenges, in turn, can alter free-running circadian period [7], and local tissue rhythms [8]. Prominent metabolic and transcriptional rhythms occur across phyla influencing physiology in cyanobacteria, fungi, plants and virtually all animals [9].

At the molecular level, the cellular clockwork involves both positive and negative transcriptional feedback loops. The BMAL1/CLOCK heterodimer is central to circadian biology, binding E-box elements (CACGTG) in the *Period* (*Per*) and *Cryptochrome (Cry)* genes and inducing their transcription [10–12]. PER and CRY proteins are shuttled to the cytoplasm forming heterodimers that interact with casein kinase Iε (CKIε). Ultimately the heterodimer translocates and returns to the nucleus where it represses BMAL1/CLOCK driven transcription [13–17]. The degradation of CRY and PER proteins relieves the repression of BMAL1/CLOCK, restarting the cycle anew [18].

Genetic, biochemical, and computational tools indicate that there is likely to be a rich network of clock modulating factors and additional clock components. Additional genetic and epigenetic mechanisms regulate tissue-specific clock functions [18–20]. RNA degradation, post-translational processing, and protein degradation are all actively regulated. Indeed, a recent study revealed that 50% of detected metabolites are under circadian control in mouse liver [21] and nearly 50% of transcripts show circadian modulation in at least one tissue [22].

Here we present data showing that Cystathionine-β-Synthase (CBS, EC 4.2.1.22), an enzyme catalyzing the first and rate limiting step of the transsulfuration pathway, regulates clock dynamics by enhancing the repressive activity of CRY1 on BMAL1/CLOCK driven transcription. Knockdown of *Cbs* by siRNA in the U2-OS and NIH3T3 cellular systems resulted in a shortened circadian period. While the period of free-running activity rhythms in transgenic mice was unchanged, the power of circadian locomotor activity rhythms was much reduced. Perhaps more strikingly, we find that CRY1 enhances CBS enzymatic activity both *in vivo* and *in vitro*. CBS enzymatic activity was significantly reduced in liver samples obtained from *Cry1*^−/−^ animals. The addition of purified CRY1 to *Cry1*^−/−^ samples restored CBS enzymatic activity. Addition of a non-binding CRY1 point mutant did not restore CBS activity. To understand the metabolic significance of CRY1 induced post-translational regulation in CBS activity, we used metabolomics to study the metabolic pathways altered in *Cry1*^−/−^ mice, utilizing *Cry2*^−/−^ and WT mice as controls. The analysis indicated that physiologic levels of CRY1 abundance specifically modulate fatty acid, amino acid and one-carbon pathways. Collectively our data show that CRY1 non-transcriptionally regulates CBS enzymatic activity and the spatio-temporal control of cellular metabolism. Additionally, wild type CBS, but not disease-causing mutant CBS-I278T enhances CRY1 repressor activity.

## Results

### CBS Physically and Functionally Interacts with the CRY1

We identified Cystathionine-β-synthase (CBS), in a computational screen for potential circadian clock components and modifiers [19]. We then employed a mammalian two-hybrid screen to identify physical interactions between CBS and a subset of known clock components. As expected, many core clock proteins physically interacted with each other, as indicated by specific activation of a pG5-*luc* reporter in transfected Human Embryonic Kidney 293T cells (HEK 293T). CBS and CRY1 were observed to interact resulting in a greater than 14-fold induction of luciferase activity (Fig.*1A* and Fig. *S1A*). No interactions were observed between CBS and the CRY1 paralogue CRY2 (Fig. *S1A* and Fig.*1A*). To identify the region of CRY1 responsible for the CBS interaction, truncated *Cry1* mutants were generated by PCR and cloned into pBIND (Fig. *S1B*). Western blot analysis of cell lysates from HEK 293T cells that expressed the truncated CRY1 constructs (from CRY-T1 to CRY1-T7) revealed expressed proteins at the expected molecular masses (Fig. *S1C*,bottom panel Western). These constructs were screened for continued interactions with CBS using the mammalian two-hybrid system [23]. As compared to full length CRY1, all of the truncated forms induced significantly less luciferase activity when co-expressed with VP16-CBS (Fig. *1A*, Fig. *S1C*). Native and Flag/His tagged co-immunoprecipitation (Co-IP) results demonstrated this interaction between CRY1 and CBS in both mouse liver and HEK293T cells (Fig. *1B* and Fig. *3B*). However, co-IP between CRY1-T1 and CBS did not show interaction (Fig. *1C*). Notably the CRY1-T1 construct retains the ability to bind with PER2 and functions as a repressor of BMAL1/CLOCK driven transcription (Fig. *S1D* and *S1E*). Hence, we concluded that the region between 586 and 606 of CRY1-T1 is required for the interaction between CRY1 and CBS.

**Fig. 1:**
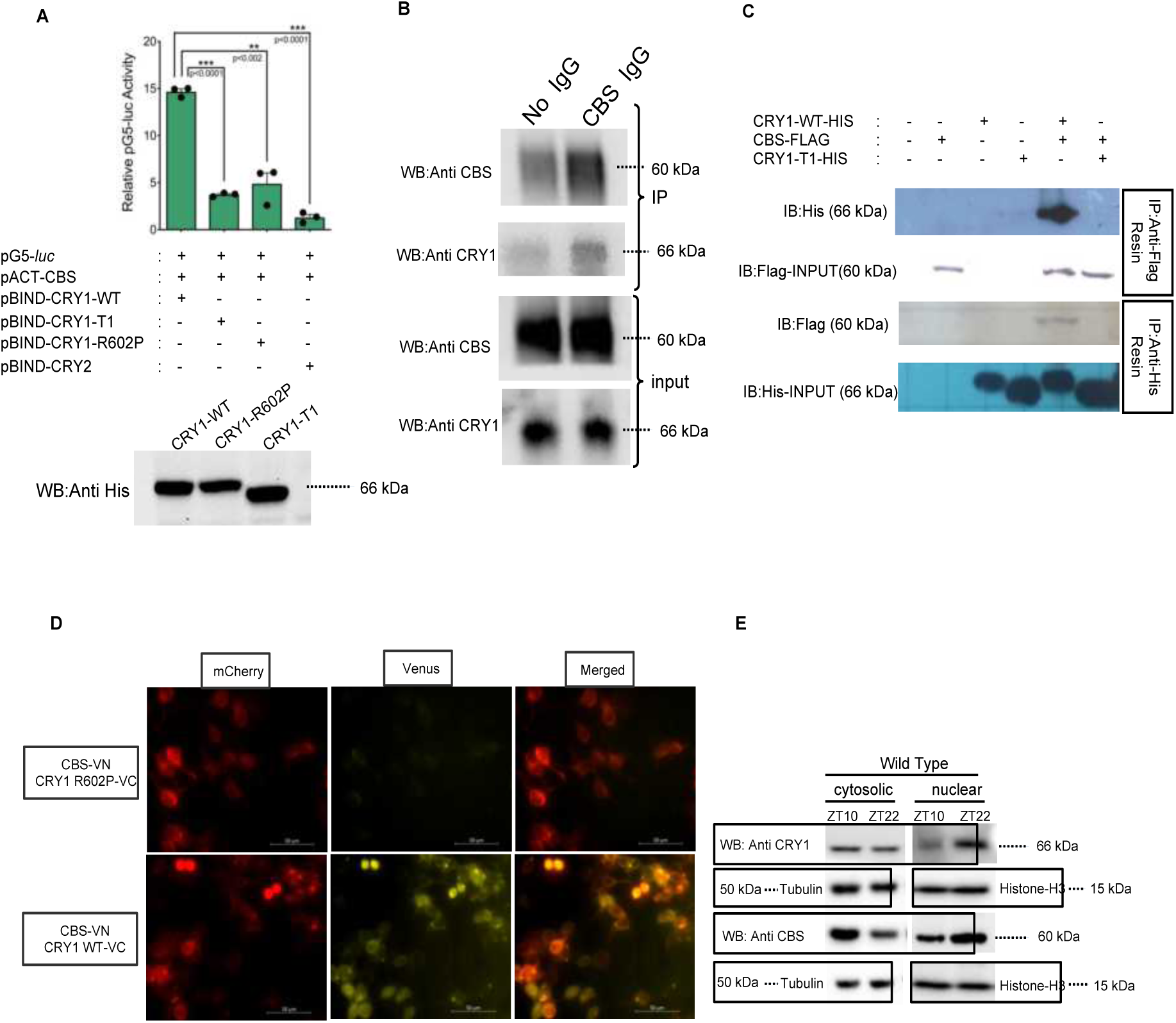
CBS physically interacts with CRY1 at both cytosol and nucleus. **(A)** Mammalian two hybrid (M2H) analysis assessing the binding of CRY1-WT, CRY1 truncation mutant (CRY1-T1), CRY2 and CRY1 point mutant (CRY1-R602P) with CBS. M2H luciferase activity is normalized with renilla luciferase and calculated relative to background. (Bars show the mean luciferase activity ± SEM, n=3, ***p<0.0001, **p<0.002) (Unpaired two-tailed t test). Western blot results indicate the expression of mutants and wild type CRYs used in M2H. **(B)** Native co-immunoprecipitation from wild type liver samples showing CRY1-CBS interaction *in vivo*. CBS and CRY1 in the input samples are shown in the bottom two panels. Upper panel shows immunoprecipitation (IP) results of CBS and CRY1. CBS antibody was pulled down with anti-mouse resin. **(C)** Co-immunoprecipitation (co-IP) experiment evaluating the binding of CBS with CRY1 and CRY1-T1 after overexpression in HEK293T cells. Either His-CRY1 or Flag-CBS was immunoprecipitated and binding partners were assessed by SDS-PAGE and Western blot with indicated antibodies. CRY1-T1 does not interact with CBS. **(D)** Bi-molecular fluorescence complementation assay (Bi-FC) of CRY1-WT and CRY1-R602P with CBS indicating both cytosolic and nuclear localizations. There are both cytosolic (mCherry overlapping) and non-overlapping venus signals. As seen from the top venus channel detection, there is no physical interaction between CRY1-R602P and CBS. All plasmids used in the assay were expressed (Fig. *S1J*). 200 ng CRY1-VC and 300 ng of CBS-VN were transfected into HEK293T cells with Fugene HD in 35-mm plates. **(E)** Fractionation of the nuclear and cytosolic components of wild type mouse liver. Fractionation efficiency is evaluated by Western blots of cytosolic (tubulin) and nuclear (Histone-H3) marker proteins (Fig. *S1O*). The relative amounts of CRY1 and CBS in both cytosolic and nuclear samples are shown (Fig. *S1P* to *S*).

The terminal 20 amino acids of CRY1 that are absent from the CRY1-T1 construct appear critical for CRY1-CBS binding. We applied PCR with degenerate primers to introduce point mutations in this region and further specify the important residues (Table S1). Constructs with randomly generated mutations were evaluated in the two-hybrid system for their interaction with CBS. Mutants yielded levels of luciferase activity, and hence CBS binding, comparable to wild type (WT). Mutant constructs from the initial screen had more than one mutation, and only constructs containing mutation at Arg602 showed reduced luciferase activity (Table S2, relative values were given in Fig. *S1F*), suggesting that CRY1 Arg602 is important for CBS binding. Queries of the NCBI Homologene database [24] revealed that Arg602 has a high degree of conservation across diverse species (Fig. *S1G*). Mutant constructs from the initial screen had more than one mutation. In order to further assess the necessity of Arg602 in CRY1-CBS interactions, we replaced Arg602 in full-length *Cry1* cDNA with proline, using site-directed mutagenesis. We confirmed expression of the mutant protein by Western blot (Fig. *1A*-below the graph). In the mammalian two-hybrid system, luciferase activity induced by the presence of CRY1-R602P and CBS was much reduced as compared to WT CRY1 and CBS. This result suggests significantly reduced interaction between CRY1-R602P and CBS (Fig. *1A*) as compared to WT control. Notably, the Arg602 to Pro mutation did not appear to interfere with either CRY1/PER2 binding or repression activity of CRY1 (Fig. *S1E*). These results suggested that CRY1-Arg602 is critically important for CBS binding.

Bi-molecular Fluorescence Complementation (Bi-FC) was performed in HEK293T to localize CRY1-CBS protein interactions within the cell. A fluorescent signal from an enhanced yellow fluorescent protein (YFP) is observed in regions where CRY1 and CBS interact. mCherry plasmid was also transfected into the same HEK293T as a cytoplasmic localization marker. DAPI was used as a nuclear localization marker. Different CBS and CRY1 interaction regions appeared in both the nuclear and cytoplasmic regions (Fig. *1D* and Fig. *S1H*; bottom merged image). We confirmed the presence of CBS in the BiFC identified interaction regions using immunohistochemistry. An anti-flag antibody demonstrated the presence of flag-tagged CBS in the interaction regions (Fig. *S1I*). While CRY1-WT interacts with CBS, CRY1-R602P does not appear to interact with CBS using the same BiFC protocol (Fig. *1D*, upper panel). Expression of CBS, CRY1, and CRY1-R602P from the Bi-FC plasmids were verified via Western blot (Fig. *S1J*).

Data from the CiraDB mouse circadian database [25], demonstrate that *Cbs* expression cycles with a 24-h period in liver and lung (Fig. *S1K, L*) [22, 25, 26]. Reconstructed human lung and liver rhythms similarly show oscillations in CBS expression [27]. Time course microarray studies demonstrate that *Cbs* expression is reduced in *Clock* mutant animals and altered circadian rhythmicity (Fig. *S1M*) [28]. We used Western blot analysis to independently confirm cycling CBS at the protein level in U2-OS cells starting from 24 hours after dexamethasone treatment to synchronize the cells (Fig. *S1N*).

### CBS and CRY1 co-localize in cytosol and nucleus *in vivo*

We reasoned that *in vivo* co-localization of CRY1 and CBS is a prerequisite for a biologically important effect. Mice were housed in a 12:12 light dark cycle. Zeitgeber time (ZT) is used to describe these environmental conditions with ZT0 being defined by lights on and ZT12 being defined by lights off. To identify the cellular distributions of these proteins, we fractionated liver samples from WT animals euthanized at ZT10 and ZT22 when the total cellular abundance of CRY1 is at its peak (ZT22) or trough (ZT10) in WT mice [29]. We separately isolated cytosolic and nuclear proteins. Proteins, known to be specifically localized in nucleus (Histone-H3) and cytoplasm (Tubulin), were used as controls to evaluate the purity of the fractions (Fig. *S1O*) [30]. As can be seen in Fig. *1E*, nuclear CRY1 abundance, like overall abundance, is higher at ZT22 as compared with ZT10 (Fig. *S1P* and Fig. *S4B*). On the other hand, cytosolic CRY1 abundance was higher at ZT10 as compared with ZT22 (Fig. *S1Q*). CBS protein was similarly observed in both the nuclear and cytoplasmic compartments. Cytosolic levels of CBS were higher at ZT10 as compared with ZT22 while the nuclear abundance of CBS was higher at ZT22 (Fig. *S1R,S*). These results further support the hypothesis that these two proteins have the potential to interact *in vivo*. We next asked if the binding of these two proteins had functional significance in circadian and/or metabolic physiology.

### Endogenous CBS Expression Modulates Circadian Oscillations

Using U2-OS and NIH 3T3 cells stably expressing a *Bmal1*-luc construct, we evaluated the influence of *Cbs* and *Cry1* on *in vitro* rhythms. A pool comprised of four individual siRNA constructs that reduced *Cbs* expression was tested alongside a pool targeted against *Cry1* and a non-targeting siRNA control (siNEG). The siRNA pools significantly (p<0.05) reduced *Cry1* and *Cbs* mRNA expression, as assessed with real-time PCR (Fig. *2C, D* - graph below the table). *Cbs* expression was reduced by 90% in NIH 3T3 cells and by ∼85% in U2-OS cells as compared to the siNEG control. Protein level knockdown was confirmed by Western blot as shown in Fig. *S2A*. Comparing the pooled results to control, *Cbs* knockdown shortened circadian period by 1.62 h and 0.69 h in NIH 3T3 and U2-OS cells respectively (p<0.05) (Fig. *2A, B*). Since CBS abundance strongly influences methionine concentration, we questioned if this affect might mediate the observed changes in period. The addition of methionine, at various concentrations, did not alter circadian period (Fig. *S2B*).

**Fig. 2:**
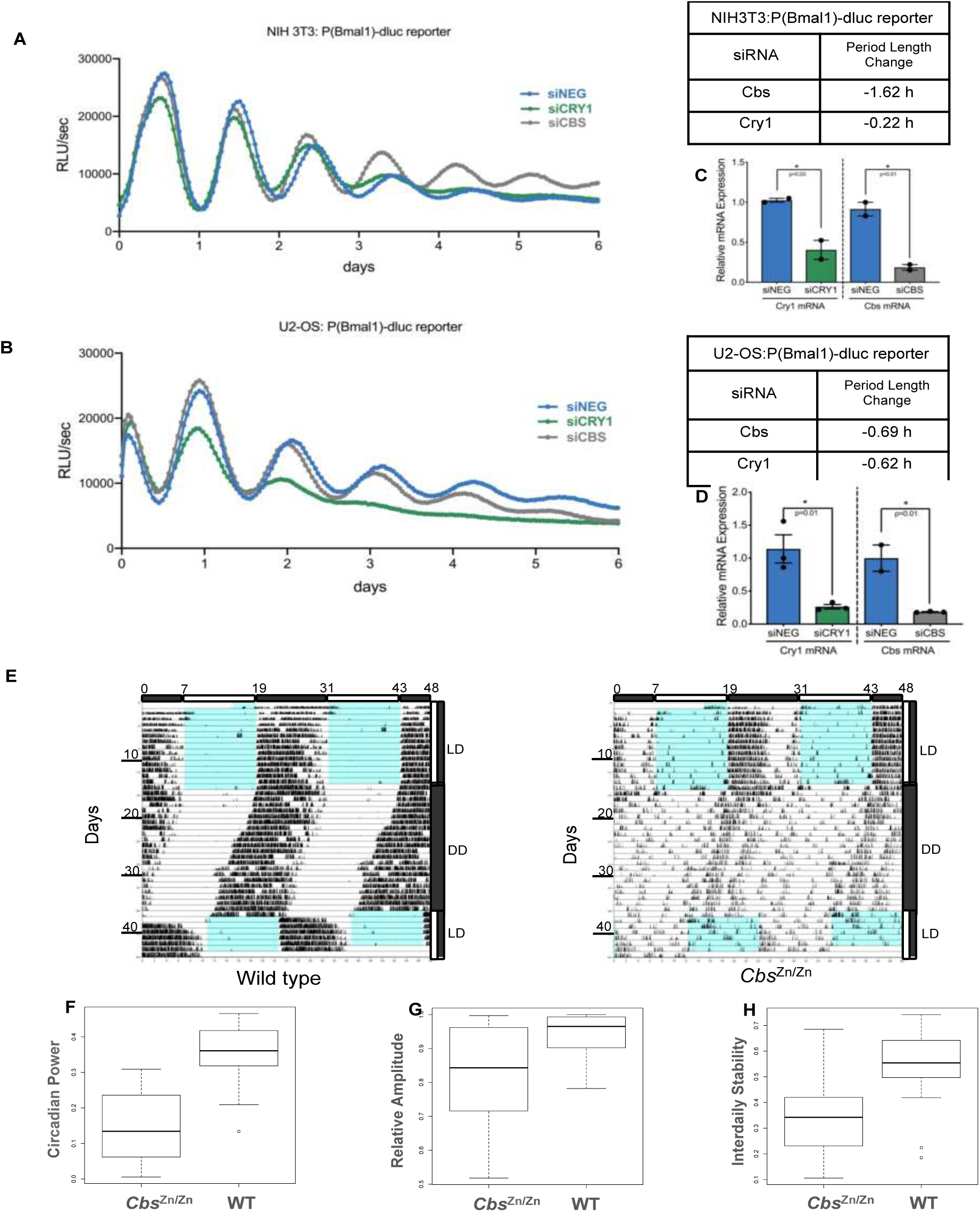
CBS influences *in vitro* and *in vivo* circadian rhythms. Bioluminescence records from **(A)** NIH 3T3:P(*Bmal1*)-d*luc* and **(B)** U2-OS:P(*Bmal1*)-d*luc* cells after siRNA mediated knockdown of the specified genes or a non-specific control (siNEG). *Cbs* knock down shortens the circadian period in NIH 3T3 (p<0.00027) and U2-OS (p<0.0017) cell lines. Knock down efficiencies in the **(C)** NIH 3T3 and **(D)** U2-OS model systems were measured by real-time PCR. RLU/sec: relative light units per second. **(E)** Representative double-plot actograms for the wheel-running activity of 6-8 weeks-old WT and *Cbs*^Zn/Zn^ animals entrained in 12:12 LD and then transferred to constant darkness. There was no statistically significant difference in net circadian period. Locomotor activity in DD was analyzed by Fourier transform. Measures of rhythm strength including **(F)** the fraction of activity power spectrum in the circadian frequency range (period ∼21-27 hours) and the non-parametric **(G)** relative amplitude and **(H)** interdaily stability indexes are plotted for both WT and *Cbs*^Zn/Zn^ animals. Circadian power (p<2.2E-10), relative amplitude (p<0.00039), and interdaily stability (p<0.000046) are all much reduced in *Cbs*^Zn/Zn^ mice. (Statistical significance assessed using a two-factor ANOVA including gender and genotype)

To examine the role of *Cbs* in modulating circadian locomotor activity, we used previously described *Cbs* transgenic mice [31]. *Cbs* is essential during murine development. To overcome this obstacle, transgenic mice that do not express murine CBS were engineered to express a hypoactive human, mutant CBS (CBS-I278T) under the control of a metallothionein promoter. Zinc driven transgene expression is able to entirely rescue the neonatal mortality [31]. After development, zinc supplementation was removed. Western blot analysis confirms the lack of both endogenous and transgenic CBS proteins in our mice [31]. The genotypes of the animals were confirmed by Western blot (Fig. *S2C*). For brevity, and despite its true genetic complexity, we designate these animals as *Cbs*^Zn/Zn^ mice throughout the manuscript.

We assessed the circadian locomotor activity of (n=26) *Cbs*^Zn/Zn^ mice along with (n=21) wild-type littermate controls. Animals were well entrained to a 12:12 light:dark (L:D) cycle for two weeks before being placed in constant darkness for 20 days. In contrast to the *in vitro* systems and in accord with experiments with CRY1^+/-^ animals, *Cbs*^Zn/Zn^ did not demonstrate in a statistically significant change in free-running circadian locomotor period. However, there was a clear change in the amplitude and stability of locomotor rhythms (Fig. *2E*). After using a Fourier Transform to construct the spectral profile of activity, we computed the fraction of the total spectral power attributed to cycles in the range of 22-26 hours. *Cbs*^Zn/Zn^ animals demonstrated much reduced circadian power. Non-parametric measures of circadian activity yielded similar results [32]. The amplitude of activity rhythms, as assessed by the difference in activity between active, and inactive periods was much reduced in *Cbs*^Zn/Zn^ mice (Fig. *2F*). Similarly, rhythm robustness, as assessed by the interdaily stability index, a measure of the similarity of activity between subsequent days, was also reduced in *Cbs*^Zn/Zn^ mice (Fig. *2G, H*). Notably these parametric and non-parametric measures are not simply measuring a reduced total level of activity in the mutant animals as they are effectively normalized by total activity levels.

### CBS modulates the circadian repressive activity of CRY1

The CBS SNP c.833T>C (p.I278T) accounts for almost 25% of all the reported *CBS* mutations found in homocystinuria patients [31, 33]. Biochemical analysis of this specific mutant indicates that CBS-I278T proteins don’t have enzymatic activity [34]. We checked if this mutant protein had the ability to interact with CRY1 using mammalian two-hybrid and co-IP tests. Although WT and mutant CBS constructs yielded comparable CBS protein levels, there was no interaction between CRY1 and CBS-I278T (Fig. *3A*). Co-IP results revealed that there was a physical interaction between WT CBS and CRY1 while no interaction was observed between CRY1 and CBS-I278T (Fig. *3B*). This result suggested that CBS-I278T could be used as negative control study performed in this section. To better understand the influence of CBS on CRY1 and circadian transcriptional function, we performed additional experiments in Neuro2A cells. HEK293T and Neuro2A cells have comparable levels of CRY1. However, Neuro2A cells demonstrate much lower endogenous CBS expression (Fig. *S3A*). *Bmal1* and *Clock* cDNA plasmids were transfected together with a plasmid containing the *Per1* promoter fused to the luciferase gene (*Per1*-d*luc*). As has been previously demonstrated, overexpression of BMAL1 and CLOCK enhanced *Per1*-d*luc* expression in unsynchronized cells. Overexpression of CRY1 or CRY1-R602P, alongside BMAL1 and CLOCK, reduced *Per1*-d*luc* activity. CBS and GAPDH cannot repress BMAL1/CLOCK driven transactivation (Fig. *S3B*). However, CBS overexpression, unlike a GAPDH control (Fig. *3C*-columns 7 and 8), enhanced the repressive activity of wild type CRY1 in a dose dependent manner (Fig. *3C*-columns 3 and 4). The addition of CBS to a system containing the otherwise functional, but non-CBS binding, CRY1-R602P mutant had no effect of luciferase activity (Fig. *3C*-columns 10 and 11). A disease-causing CBS-I278T mutant, which doesn’t interact with CRY1, did not affect the repressive activity of CRY1 (Fig. *3C* columns 5 and 6). The expressions of wild type and mutant CBS were assessed by Western blot (Fig. *S3C*). Taken together, these data demonstrate that CBS-CRY1 binding enhances the repressive activity of the CRY1 on BMAL1/CLOCK driven transcription. We hypothesized that decreased CBS protein abundance would reduce CRY1 mediated repression and increase the expression of direct clock-controlled output genes. We first tested this hypothesis in the U2-OS model system, assessing the expression of the canonical E-BOX output genes *Dbp* and *Per2* after siRNA mediated *Cbs* knockdown. As shown in Fig. *3D*, knocking down *Cbs* to 10% basal levels increased the expression of both *Per2* and *Dbp*. We then confirmed these results *in vivo,* comparing the expression of *Per2* and *Dbp* in the livers of WT and *Cbs*^Zn/Zn^ mice sacrificed at ZT12 (when CBS levels peak*). Dbp* and *Per2* expression levels were significantly higher in *Cbs*^Zn/Zn^ mice (Fig. *3E*). Notably CBS abundance did not significantly influence the expression of CRY1 itself (Fig. *3E*-middle panel) in either experiment, suggesting that the CBS exerts these effects post-translationally through altered CRY1 repressive activity.

**Fig 3:**
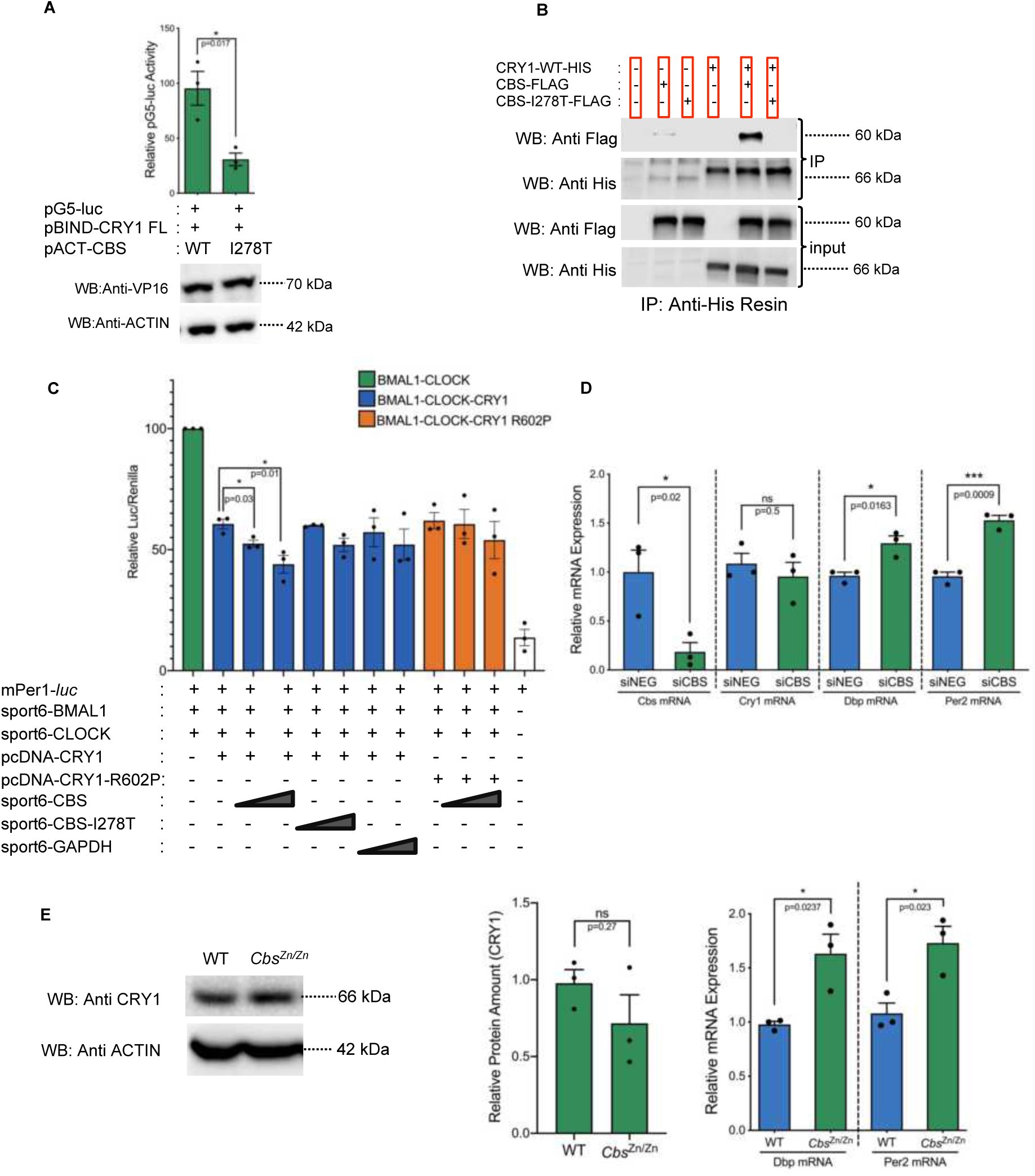
Wild type, but not mutant CBS, enhances CRY1 repressor activity. **(A)** Mammalian two-hybrid assay assessing binding between CRY1 and disease-causing mutant CBS (CBS-I278T). The CBS-I278T point mutant significantly reduces the interaction with CRY1 (n=3, mean ± SEM, p=0.017, unpaired two-tailed t-test). The blot below shows the levels of CBS WT, CBS mutant, and Actin expression. **(B)** Co-immunoprecipitation showing the loss of physical interaction between CRY1 and CBS-I278T. Indicated proteins were overexpressed in HEK293T and His tagged CRY1 was pulled down with Anti-His resin. CRY1 and CBS were detected with Anti-His and Anti-Flag antibodies in both input and immunoprecipitation (IP) samples. **(C)** A Per1:dluc reporter assay was used to assess the influence of selected proteins on BMAL1-CLOCK transcriptional activity. Luciferase expression was normalized by Renilla luciferase expression. (n=3, mean ± SEM, *p=0.01, * p=0.03) (unpaired two-tailed t test). **(D)** The effect of siRNA mediated CBS knockdown on the expression of selected transcripts in U2-OS cells. Relative mRNA expressions were assessed by real-time PCR. **(E)** Expression of the selected transcripts in the livers of WT and *Cbs*^Zn/Zn^ animals euthanized at ZT12 and assessed by real-time PCR. CBS deficiency increased the transcription of Dbp and Per2 without altering the Cry1 mRNA or CRY1 protein levels both *in vitro* and *in vivo*.

### CRY1 regulates the enzymatic activity of CBS

In order to investigate the reciprocal influence of CRY1 on CBS enzymatic activity we employed a well-established colorimetric assay based on the production of H2S [35, 36]. Cystathionine-γ-lyase (CSE) also contributes to H2S production in liver. In order to directly assess the influence of CRY1 on CBS activity, we inhibited the activity of CSE using PAG (DL-propargylglycine) as has been previously described [37]. Despite a background level methylene blue production, the measured enzymatic activity in liver lysate obtained from *Cbs*^Zn/Zn^ animals was significantly reduced as compared to the lysate from WT animals (Fig. *S4A*). Wild type, *Cry1^−/−^* and *Cry2^−/−^* C57Bl6/J mice were euthanized at either ZT22 or ZT10, when the total cellular abundance of CRY1 is at its peak or trough, respectively [17, 38] (Fig. *S4B*). Mouse genotypes were confirmed before starting experiments (Fig. *S4B*, C). Consistent with previous work [39], our data demonstrate that the amount of the CRY1 in the livers of *Cry2^−/−^* animals is greater than wild type (Fig. *S4D, E*), while CRY1 oscillation is still maintained (Fig. *S4F*). Accordingly, we studied *Cry1^−/−^* animals using both WT and *Cry2^−/−^* mice as controls.

We performed Western blots using cell-free protein extracts from WT livers obtained at ZT10 and ZT22. Consistent with the known oscillation in *Cbs* expression, there was more CBS protein at ZT10 than at ZT22 (Fig. *S4G*). The total CBS enzymatic activity was correspondingly higher at ZT10 (Fig. *S4H*).

There was no statistically significant difference in CBS abundance when comparing the livers of WT, *Cry1^−/−^* and *Cry2^−/−^* animals, euthanized at the same time. (Fig. *S4I, J*). Despite the comparable amounts of CBS at ZT10 across these genotypes, CBS activity was significantly lower in extracts from *Cry1^−/−^* animals as compared to WT (Fig. *4A*). Similarly, CBS activity was higher in samples from *Cry2^−/−^* animals as compared to WT, consistent with the compensatory increase in CRY1 (Fig. *4A, B* bottom panels, quantification is in Fig. *S4 D,E*). Thus, CRY1 concentration appears to correlate with CBS enzymatic activity *in vivo*.

**Fig. 4:**
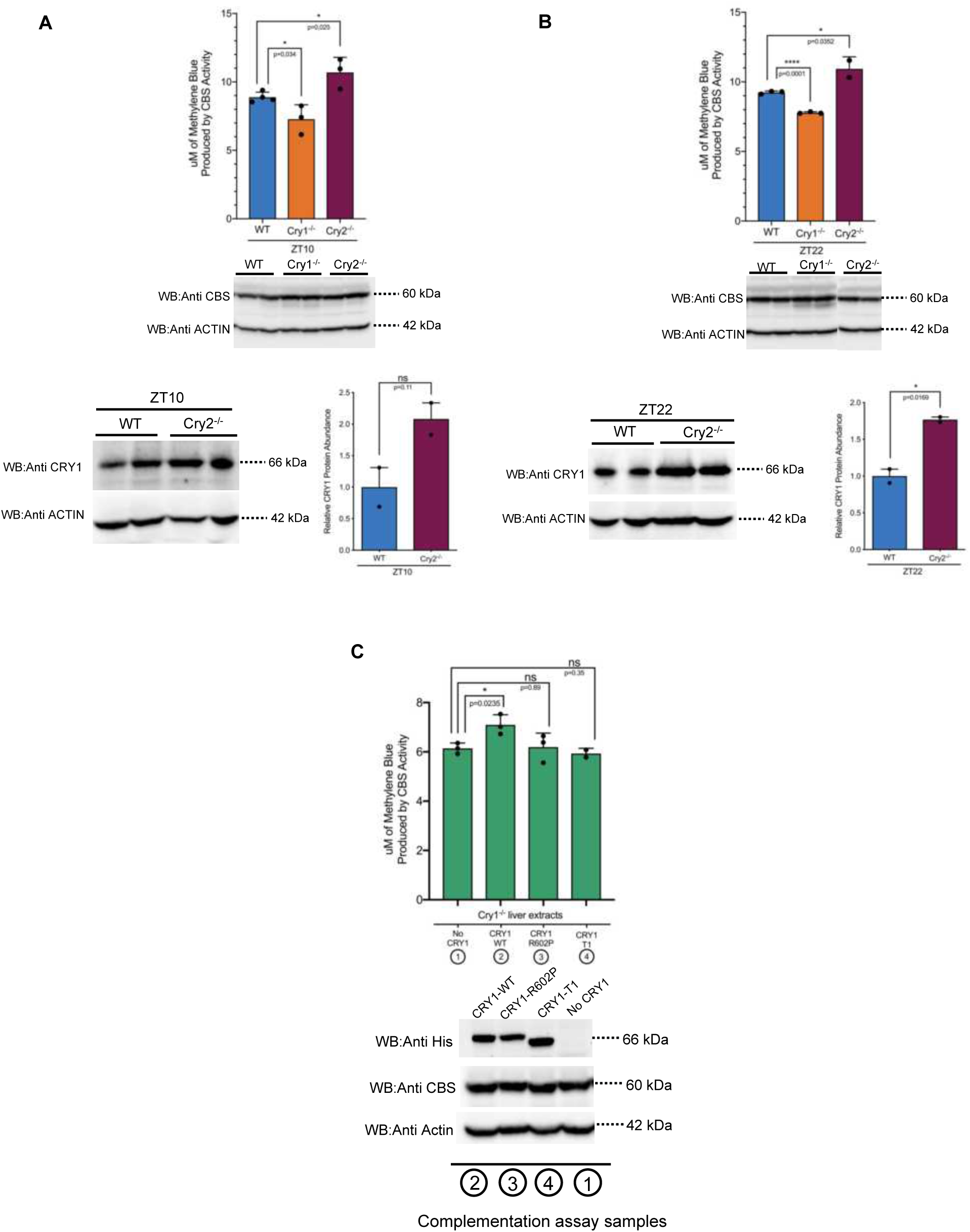
CRY1 enhances CBS enzymatic activity. **(A) *Top:*** CBS enzymatic activity in wild type, *Cry1*^−/−^and *Cry2*^−/−^mouse liver samples obtained at ZT10 (n=at least 3 biological replicates, mean ± SEM, *p=0.034, *p=0.025). Enzymatic activity is assessed by a colorimetric H2S production assay. ***Middle:*** Western blots showing that samples were diluted to equalize CBS protein concentrations. (n=2 biological replicates). ***Bottom***: Western blot indicating increased CRY1 in *Cry2*^−/−^ samples. **(B)** The same data is shown for WT, *Cry1*^−/−^, and*Cry2*^−/−^mouse liver samples obtained at ZT22. **(C)** Complementation assay showing that the addition of exogenous CRY1 WT restores reduced CBS activity in *Cry1*^−/−^liver samples. Non-binding CRY1-T1 or CRY1-R602P mutant proteins do not complement. The Western blot in the right shows the expression level of CBS, mutant and wild type CRYs used in this experiment.

Assay results for ZT22 mice paralleled the results from ZT10 mice (Fig. *4B*). In summary, liver lysate samples with absent CRY1 demonstrated reduced CBS activity, while samples with increased abundance of CRY1 demonstrated augmented CBS enzymatic activity.

We questioned if the lack of CRY1-CBS protein binding is sufficient to explain the observed reduction of CBS activity in *Cry1^−/−^* liver lysates. We tested if the addition of exogenously overexpressed CRY1 protein would restore CBS activity. Plasmids expressing wild type *Cry1*, *Cry1*-R602P and *Cry1*-T1 cDNA were transfected to HEK293T cells. Crude extracts were prepared from each sample and the same amount of the mutant or wild type CRY1 proteins were used in each assay (Fig. *4C*, bottom panel). No significant differences were noted in CBS protein abundance (Fig. *S4K*). Addition of extract containing wild type CRY1 protein to *Cry1^−/−^* liver extracts increased the relative enzymatic activity of the CBS. The addition of extracts of either CRY1-R602P or CRY1-T1 mutant proteins (both are unable to interact with CBS) did not restore CBS activity to wild type (Fig. *4C*).

Collectively these data support the theory that the physiologic binding of CRY1 to CBS enhances CBS enzymatic activity both *in vivo* and *in vitro*.

### Metabolomic studies support the importance of CBS-CRY1 interactions *in vivo*

CBS is a rate-limiting, regulatory branch point in the eukaryotic methionine cycle. CBS catalyzes a pyridoxal 5′-phosphate dependent beta-replacement reaction condensing serine and homocysteine to cystathionine. Cystathionine is subsequently converted to cysteine in a reaction catalyzed by cystathionine γ-lyase. Meanwhile, homocysteine, an intermediary amino acid metabolite in this process, is critical for the regulation of methionine, folate, and transsulfuration pathways [40]. The absence of the CBS in mice results in high levels of total homocysteine and a reduced SAM/SAH ratio in serum [41].

We hypothesized that the temporal modulation of CBS enzymatic activity, through its dynamic interaction with CRY1 might influence these critical pathways. In order to compare the changes resulting from *Cbs* and *Cry* knockouts, we performed two independent, metabolomic studies. We first identified metabolites with significant differences in hepatic abundance comparing *Cbs*^Zn/Zn^ mice and WT littermate controls (n=23, ∼5-6 mice of each gender and genotype) sacrificed at ZT12. The influence of CBS genotype was far-reaching. A two factor ANOVA including gender and genotype identified 81/194 metabolites significantly modulated by CBS genotype (> FDR of 5%) (File *S1*). KEGG pathway analysis, implemented with Metaboanalyst, demonstrates that homocysteine and methionine pathway components were strongly overrepresented among the CBS affected metabolites (Fig. *5A*-vertical axis and File *S2*). Moreover, the affected metabolites played a central role in this pathway as assessed by network centrality (Fig. *5A*-horizontal axis).

**Fig. 5:**
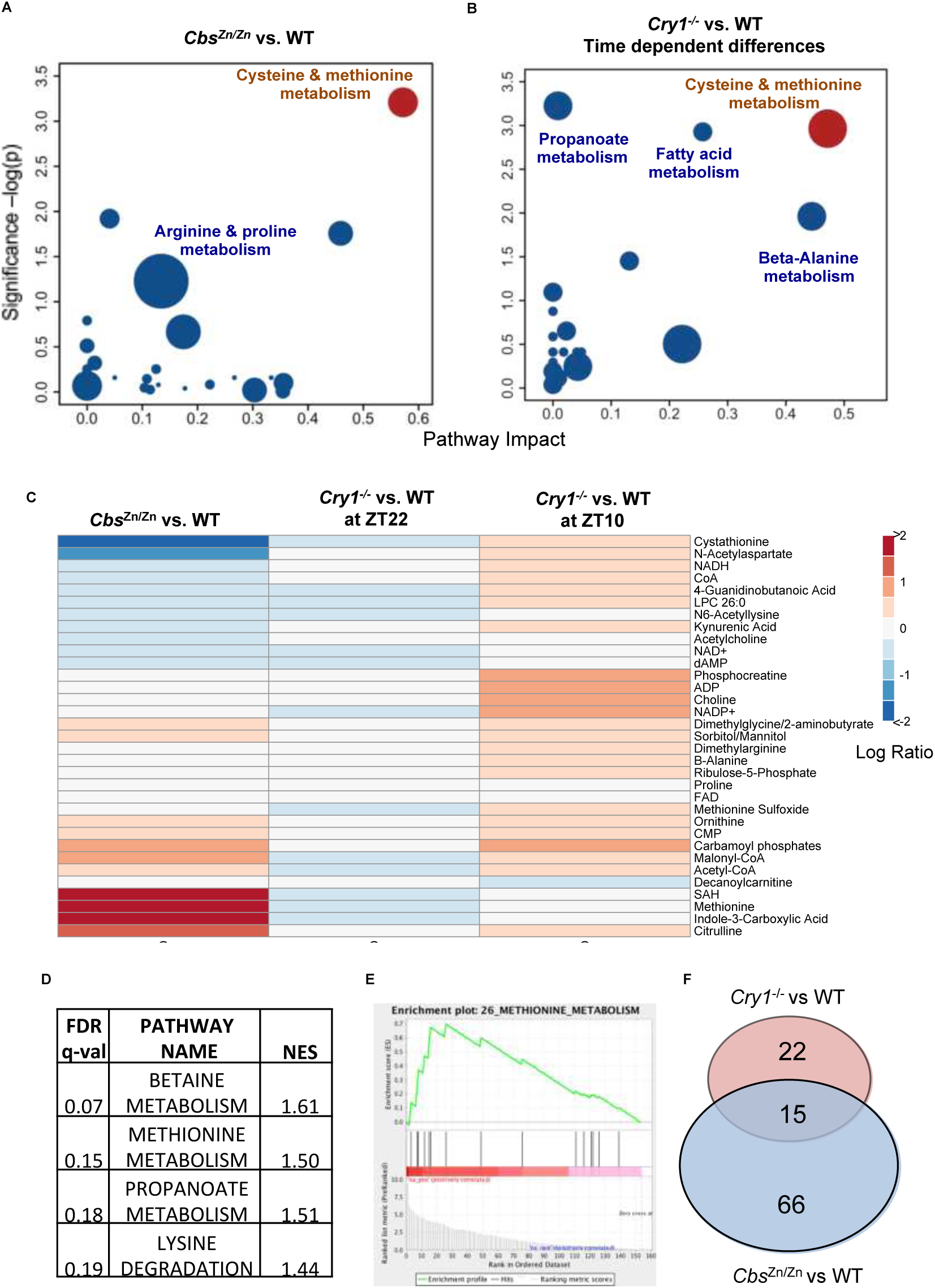
Metabolomic analysis of CBS and Cryptochrome Mutants. **(A)** Pathway overrepresentation describing metabolomic changes in the liver of *Cbs*^Zn/Zn^ animals. Metabolites differentially abundant (FDR <5%) between *Cbs*^Zn/Zn^ and WT littermate controls were compared to the full list of tested metabolites. KEGG pathways that are overrepresented among CBS affected metabolites are shown. The vertical axis shows the significance of overrepresentation. The horizontal axis depicts the importance of the CBS affected metabolites in the pathway as assessed by network centrality. Cysteine and methionine metabolism are strongly affected by the CBS deficiency. **(B)** Pathway overrepresentation describing liver metabolites influenced by the interaction of *Cry1^−/−^*genotype and circadian time. Metabolites modulated by the *interaction* between cryptochrome genotype (*Cry1^−/−^* vs WT) and circadian time (ZT10 vs ZT22) were identified by ANOVA (p <0.05). Cysteine and methionine metabolism are strongly affected by the interaction, consistent with the hypothesis that CRY1 binds CBS and dynamically alters its enzymatic efficiency. **(C)** Heat map comparing the metabolomic influence of CBS deficiency (in *Cbs*^Zn/Zn^ mice) with the dynamic metabolomic influence of CRY1 genotype. Metabolites significantly affected by the interaction between CRY1 genotype and time are shown. At ZT22, when CRY1 expression is highest in WT animals, the metabolite differences between WT and *Cry1*^−/−^mice appeared similar to the differences between WT and *Cbs*^Zn/Zn^ mice. **(D)** Set enrichment analysis was performed on the full list of measured metabolites, ranked by the statistical significance of the interaction between time and Cryptochrome genotype. Annotated metabolite sets were obtained from the Small Molecule Pathway Database. The statistical significance of enrichment and the Normalized Enrichment Score (NES) for each set is shown. **(E)** Methionine related metabolites are highly enriched near the top of the list of those modulated by the time genotype interaction. **(F)** Venn diagram showing overlap of individual metabolites significantly modulated in the two experiments. The blue circle depicts metabolites significantly affected by CBS genotype. Pink depicts metabolites influenced by the interaction between circadian time and Cryptochrome genotype.

We next compared liver samples from *Cry1*^−/−^ and WT mice (n=5 or 6 of each gender and genotype) sacrificed at either Z10 or ZT22 (Fig. *5B*). A preliminary analysis focused on the role of gender in influencing metabolite abundance in the WT mice. WT mice of both genders were also used in the *Cbs*^Zn/Zn^ experiment. This analysis allowed us to evaluate the internal consistency of our own experimental results and the consistency or our results with published literature. We computed the log ratio of mean metabolite abundance in female as compared to male WT mice for both experiments. Metabolites highly abundant in male or female WT mice in one experiment showed a similar trend in the other experiment (Fig. *S5A*). Based on separate ANOVA analyses of these two experiments, 30 metabolites were identified as being significantly influence by gender in the control animals for the *Cry1*^−/−^ experiment while 29 were identified in the control animals from the *Cbs*^Zn/Zn^ experiment (FDR <15%) (Fig. S5B and File *S1*). The overlapping 11 metabolites behaved very similarly in both datasets (Fig. *S5C*). Notably many of these metabolites, including carnitine derivatives, have been identified as showing sexually dimorphic abundance in several tissues including liver [42]. Ranking the full list of metabolites based on the relative abundance in female as compared to male mice and performing set enrichment analysis [43] (Fig. *S5D*). We find that glycolysis and gluconeogenesis are modulated by gender at a pathway level, a finding that has been observed by others [44]. These control data also revealed metabolites with different concentrations in WT animals at ZT10 vs. ZT22. Nine of the 18 “core” or “multi cycling metabolites” identified by Krishnaiah et al [21] showed significant, time modulated differences in abundance (p <0.05, ANOVA including time and gender). These metabolites include L-histidine, Glutathione, phosophcholine, UDP-glucose, methionine, SAH, cystathionine, and 1-methylnicotinamide (File *S2*). Set enrichment analysis (Fig. *S5E*) demonstrated differences in pathways relating to purine, pyrimidine, and glutamate metabolism consistent with earlier findings [45].

Returning to the circadian modulation of CBS activity, we reasoned that CBS/CRY1 interactions depend on CRY1 abundance and hence circadian time. Hence, we sought to identify metabolites with abundances that differ between the genotypes but in a manner that depends on time. We used a 3-factor ANOVA that included time, genotype, and gender (File *S3*) to identify metabolites modulated by the interaction between circadian time and cryptochrome genotype (34 of 178 measured metabolites were identified at a p-value <0.05 while 15 were identified at an FDR <15%). Consistent with our hypothesis that these changes would be mediated by the presence or absence of CBS interactions, pathway overrepresentation analysis (File S4) demonstrates that the most prominent changes occurred in homocysteine and methionine metabolism (Fig. *5B*). The heatmap shown in Fig. *5C* further supports this relationship. At ZT22, when CRY1 expression is highest in WT animals, the metabolite differences between WT and *Cry1*^−/−^ mice appeared similar to the differences between WT and *Cbs*^Zn/Zn^ mice. On the other hand, at ZT10 when CRY1 expression is near its nadir, the differences between *Cry1*^−/−^ and WT animals in were muted and in a few cases reverse direction. For example, the levels of the cystathionine and NADP+ are lower in *Cbs*^Zn/Zn^ mice as compared to controls. When comparing *Cry1*^−/−^ mice and controls, this same relationship is observed at ZT22 but not ZT10.

The metabolites significantly influenced by the interaction between CRY1 genotype and circadian time demonstrated a very consistent response among male and female mice. While some of these metabolites showed a trend for higher (mean) expression in one gender or another, when abundances are independently normalized for male and female mice, parallel responses to time and genotype are evident in mice of both sexes (Fig. *S6A*).

The choice of numerical cutoffs (e.g p<0.05) can greatly influence overrepresentation results [43]. We used metabolite set enrichment analysis using the full list of tested metabolites ranked by the significance of the genotype-time interaction. Methionine metabolism was among the pathways demonstrating the strongest enrichment (Fig. *5D, E*). Broadening our search to include smaller pathways demonstrates that the interaction of time and *Cry1* genotype also influences fatty acid metabolism (Fig. *5B*).

We then defined a custom set of CBS sensitive metabolites using the results of the *Cbs*^Zn/Zn^ vs WT metabolomic study. Based on our ANOVA analysis (File *S1*) of these data, we created a set including all metabolites significantly affected by CBS genotype (FDR < 10%). These CBS sensitive metabolites are significantly enriched among metabolites modulated by the interaction of *Cry1* genotype and time (p= 0.05, Normalized Enrichment Score=1.23, Fig. *S6B*).

Finally, we compared the hepatic metabolomic profile of the female *Cry1*^−/−^ mice euthanized at ZT10 and ZT22 with the profiles observed from an available group of female *Cry2*^−/−^ mice euthanized at these same times. ANOVA was again used to identify metabolites modulated by the interaction of genotype and time. Cystathionine metabolism was again the most strongly influenced pathway (Fig. *S6C*, Files *S5,6*).

In each of these metabolomic comparisons, differences were also noted in metabolites and pathways closely connected to homocysteine metabolism including lysine, arginine and proline metabolism, (Fig. *5A,B*, *S6B* and *S7A*) suggesting more distributed changes in amino acid metabolism.

Homocysteine occupies a critical metabolic branchpoint: it can be either converted into cysteine by transsulfuration or re-methylated back into methionine using either by methyltetrahydrofolate or betaine as methyl donors [37]. CBS induced changes in homocysteine levels are thus predicted to affect both the methyl cycle and collateral pathways. Many metabolites influenced by the interaction of time and cryptochrome genotype reflect these peripheral changes (Fig. *S7A,B*). As seen from Fig. *S7A*, we observed that circadian differences in several metabolites involved in homocysteine metabolism were blunted in *Cry1*^−/−^ mice. Many of these same metabolites demonstrated augmented circadian differences in *Cry2*^−/−^ mice (Fig. *S7A*), consistent with a compensatory increase in CRY1.

Hepatic SAH was significantly affected by the interaction of circadian time and genotypes reflecting changes in cellular methylation potential. Citrulline and ornithine, components of urea cycle and metabolites important to arginine and proline metabolism were also affected (Fig. *S7A*). Choline metabolism is closely connected with the CBS metabolism as the conversion of betaine is coupled with conversion of homocysteine to methionine [40]. Indeed, betaine metabolism was the pathway most significantly enriched for metabolites influenced by the interaction between time and CRY1 genotype (Fig. *5D*). Acetylcholine, choline and dimethylglycine were all modulated by the interaction of time and Cryptochrome genotypes (Fig. *5* and Fig. *S7*). *Cry1^−/−^* and *Cry2^−/−^* mice have been previously found to have altered fatty acid metabolism [46]. We found that cryptochrome knockouts have altered time dependent levels of malonyl CoA, Acetyl CoA and NADP+ (Fig. *5* and Fig. *S7*). These metabolites are essential components of fatty acid metabolism and modulated by cellular redox potential. Analysis of wild type and *Cbs*^Zn/Zn^ liver samples revealed the changes in the concentrations of these same metabolites suggesting that the interaction between CRY1 and CBS regulates the methyl and fatty acid metabolisms at physiological level. Unlike homocysteine metabolism, however, the effects of CRY and CRY2 deficiency on the fatty acid related metabolites appeared to be similar. This may reflect the limited temporal resolution of our data or a distinct mechanism of regulation.

## Discussion

In addition to controlling locomotor behavior, the clock adjusts metabolic pathways in anticipation of daily changes in nutrient flux and energy needs [22]. Recent studies have highlighted the bi-directional regulation between circadian rhythms and metabolic pathways in mammals [8]. The molecular mechanism(s) underlying this regulation are, however, not well understood. In this study, we provide direct evidence that CRY1 and CBS interact both physically and functionally, contributing to the transcriptional regulation of clock-controlled genes, and more strikingly the post translational regulation of methyl cycle and amino acid metabolisms.

Proteins in the Photolyase/Cryptochrome family perform diverse functions in various organisms [47]. Sequence alignment analysis shows that CRYs have variable extended C-terminal domains that range from 30 to 300 amino acids [48]. The Intrinsically Disordered Region (IDR) of the C-terminal domain enables cryptochrome proteins to interact with various proteins, facilitating tissue and organism specific functions [48]. Our study indicates that Arg602 in the IDR region of CRY1 is essential for the interaction with CBS (Fig. *1*). Further experiments showed that this interaction modulates the activity of both proteins. CBS enhances CRY1 repression activity while CRY1 in turn enhances CBS enzymatic activity (Fig. *3* and Fig. *4*). The effect of CBS on CRY1 activity is likely to be important for the control of circadian period at the cellular level (Fig. *2A, B*) and may play a role in diet induced circadian alterations in the liver. While developmental effects cannot be excluded, CBS appears to contribute to the strength and robustness of *in vivo* circadian locomotor activity without affecting the period (Fig. *2C*). Several previously identified clock gene modifiers and mutants have different or more extreme effects on *in vitro* rhythms as compared to *in vivo* locomotor behavior [5, 49]. In particular our *in vitro* analysis suggested that *Cbs* deficiency reduces but does not nearly eliminate CRY1 mediated repression. *Cry1* knockdown reduces *in vitro* period but heterozygote Cry1^−/+^ mice do not demonstrate a circadian impairment in behavior [50].

Our metabolomics results highlighted the physiologic importance of this interaction. The reduction of CRY1 alters one carbon and amino acid metabolism, metabolic reactions that are central to life. The core circadian proteins CRY1, CRY2, PER1, PER2, BMAL1, and CLOCK all bind the CBS gene with CRY1 specifically binding its promoter [51]. A recent study demonstrated that the accessory circadian protein NR1D1 also binds the CBS promoter [52]. Nascent-seq and other studies highlight the importance of active, circadian regulation of CBS mRNA degradation [53, 54]. Thus, the post-translational regulation of CBS enzymatic activity by CRY1 binding provides yet another level of regulation. We speculate that this non-transcriptional influence of CRY1 on CBS activity has several key advantages. The circadian production and degradation of proteins is energetically costly. As the increases and decreases in CBS and CRY1 abundance are, well synchronized in the cytoplasm (Fig. *S1Q,R*), CRY1 can increase enzymatic efficiency while moderating the protein synthesis burden. Alternatively, in the nucleus, differently timed interactions may blunt CBS induced changes in redox potential and SAM availability, while allowing metabolic feedback on transcriptional circadian rhythms. We suspect that multi-leveled circadian control of CBS reflects the importance of methyl cycle and amino acid metabolism and cellular physiology. This regulatory role of CRY1 in modulating CBS activity adds to its known ability to bind and modulate G-proteins [55].

CBS and CRY1 demonstrate temporally and spatially dynamic cellular distributions. Our initial analysis focused on ZT22 and ZT10 because these are the times of peak and trough CRY1 abundance in whole cell lysates. The abundances of CRY1 and CBS are anti-phasic in whole cell lysates. However, the bulk of CRY1 is localized to the nucleus and the bulk of CBS is localized in cytoplasm. While we only measured two time points, focusing specifically on cytoplasmic CRY1 and CBS levels the two proteins appear to be in-phase (Fig. *S1Q,R*). A full understanding of this biology will likely require a more detailed time course analysis separately evaluating nuclear and cytoplasmic fractions. Indeed, we hypothesize that the sometimes-modest metabolic effects we observed may underestimate larger difference likely observed following peak and trough cytoplasmic levels.

While we have focused on the role of CBS in the synthesis of cystathionine, in the last decades the role of CBS in the synthesis of H2S has received increased attention. Indeed H2S production was directly assessed in the colorimetric assays used in the enzymatic studies above as it utilizes same CBS domains and binding sites. H2S, a gaseous transmitter, plays an important role in many physiological processes [56]. Three enzymes can independently catalyze the formation of H2S: (1) CBS, (2) cystathionine-γ-lyase (CSE) and (3) 3-mercaptosulfurtransferase (3-MST). CBS is present in highest abundance in the central nervous system, liver and pancreas. CSE is primarily responsible for H2S production in the cardiovascular system [57]. 3-MST is located predominantly in the mitochondria and produces H2S in concert with cysteine aminotransferase. Thus, we hypothesize that the CRY1 might directly influence H2S production in brain and liver through its interaction with CBS. H2S signaling is important in maintaining proper CNS function [58]. Dysregulated H2S has been implicated in neurodegenerative diseases, including Alzheimer’s disease [59]. Our data suggest the possibility that CNS H2S production is post-transcriptionally modulated by the CRY1 and molecular circadian clock. Disturbed clock function is often a hallmark of neurodegeneration. We can only speculate if disturbed circadian rhythms in H2S production contribute to, or are a consequence of, neurodegenerative diseases. Of equal clinical interest, the relationship between H2S, CBS function, and cancer has been the subject of several recent investigations [60]. CBS mediated overproduction of H2S in colorectal and ovarian cancer cells enhances cell proliferation, migration and cellular bioenergetics. As CRY1 enhances CBS activity and H2S production, CRY1-CBS interactions could be a potential new chronotherapeutic target.

Studies with clock-disturbed mice provide increasing evidence for the importance of intact circadian rhythms for healthy metabolism. A recent study showed that 50% of evaluated metabolites exhibit a circadian rhythm in liver [21]. Many metabolic signaling molecules also demonstrate robust circadian regulation [61]. For example, the circadian clock is required for the secretion of insulin and proper development of the beta cells in the pancreas [62]. Metabolic feedback on the regulation of core clock proteins through has been described, including cross-talk between core clock proteins, nuclear receptors [63], and histone deacetylases [64, 65]. Recently, it has been shown that CLOCK directly acetylates argininosuccinate synthase to regulate arginine biosynthesis [66]. Here, we provide evidence that a new mode of bi-directional regulation, where CRY1 directly interacts with a key enzyme, CBS, enhancing its activity and regulating one-carbon and urea metabolism. Metabolomic analysis of liver samples obtained from *Cry1*^−/−^, *Cry2*^−/−^, and WT mice allowed us to demonstrate the significance of this dynamic interaction at physiological abundances (Fig. *6*).

**Fig. 6:**
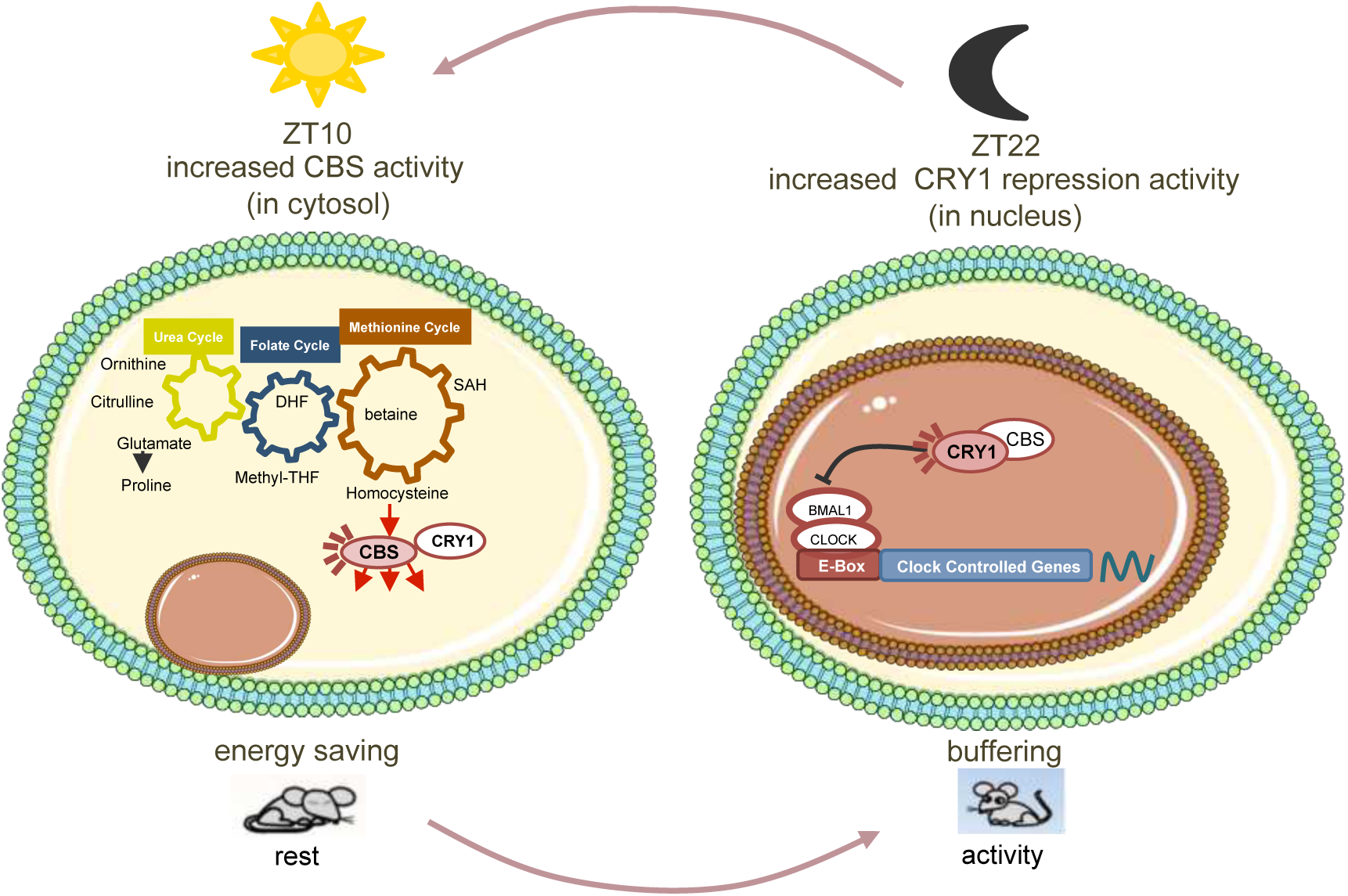
Model of CRY1-CBS interactions directly linking circadian and metabolic functions. When cytosolic CRY1 abundance is high (ZT 10 compared to ZT 22) CBS enzymatic activity is enhanced. The energetic costs that would be required to synthesize new CBS enzyme are reduced. When nuclear CBS concentration is higher (ZT22 as compared to ZT10) CBS enhance the repressive activity of CRY1 in local tissue clocks.

Notably CRY1 does not appear to interact with the clinically important CBS-I278T mutant (Fig. *3A, B*). CBS I278T is the cause of 25% of all CBS related homocystinuria disorders [31]. Our data raises the possibility that CBS I278T mutation also disrupts the circadian modulation of cysteine and amino acid metabolism. Timed administration of folate which has short serum half-life may help restore the metabolic rhythms. The clinical importance of these rhythms will need to be assessed.

In conclusion, our findings indicate that physiologic CBS-CRY1 binding influences circadian physiology and metabolism. CBS-CRY1 interactions provide a post-translational switch regulating the spatio-temporal activity of CBS and CRY1. In the cytosol, CRY1 and CBS abundances both peak near ZT10 when the enhancement of CBS activity has the greatest potential affect. In the nucleus, CBS likely acts at a different time modulating the repressive activity of CRY1 on BMAL: CLOCK-driven transcription (Fig. *6*). While the details of the interaction will need to be more thoroughly studied, this work highlights the importance of post-translational mechanisms in coupling central metabolic pathways with circadian control. It also opens up avenues for the study and treatment of CBS related disease.

## Materials and Methods

### Reporter Assays with Transient Transfection

#### Plasmid Constructs

*Bmal1, Clock, Per1, Per2, Per3, Cry1, Cry1-T1, Cry1-R602P* and *Cry2* were subcloned into the pBIND vector and *Cbs* and *Per2* were subcloned into the pACT vector for Mammalian-Two-Hybrid experiments according to the instructions of Promega CheckMate™ Mammalian Two-Hybrid System. Luciferase gene pG5-*luc* was used as the reporter gene, while pGL4-*Renilla* was used for normalization of transfection efficiencies. *Cbs*, *Cry1*, *Cry1-T1*, *Cry1-R602P, Bmal1* and *Clock* were subcloned into pCMV-sport6 vector separately for repression assay. An mPer1-*luc* construct was used in the transfection based-reporter assay.

#### Mammalian-two-hybrid

For mammalian two-hybrid assays, 25 ng of pGL5, 5 ng Renilla luciferase, 50 ng pACT, and 50 ng pBIND plasmids were transfected into HEK 293T cells in 96-well plates as previously described [23]. Transfected cells were analyzed after 30 h incubation for luciferase reporter activity with DualGlo luciferin reagent (Promega) using a luminometer (Thermo Fluoroskan Ascent FL). The firefly luciferase activity was normalized using *Renilla* luciferase for transfection efficiency.

#### Per1-Luc Reporter Gene Assay

Neuro2A (N2A) cells were grown in DMEM with 10%FBS. The day before transfection, cells were seeded into 10-cm dishes to have 80% confluency on the next day. Cells were transfected with Fugene HD (Promega) transfection reagent with the indicated amount of plasmids: 50 ng pCMV-sport6*-Bmal1*, 125 ng pCMV-sport6-*Clock*, 50 ng *mPer1-luc*, 5 ng pGL4-*Renilla*, 3 ng pcDNA-*Cry1* (or pcDNA*-Cry1-R602P*), 20 or 30 ng pCMV-sport6-*Cbs, or pCMV-sport6-Cbs I278T, or pCMV-sport6-GAPDH* and empty pCMV-sport6 in order to bring the total DNA amount to 300 ng. The preparation of cell lysates and dual luciferase assay were carried out according to the manufacturer’s protocol (Promega). After 24 h, activity of both firefly and *Renilla* luciferase was measured using a luminometer (Thermo Fluoroskan Ascent FL). The firefly luciferase activity was normalized using *Renilla* luciferase for transfection efficiency. All experimental results are the average of three independent experiments.

#### Cell Based Circadian Assay

Either U2-OS or NIH3T3 cells were plated on 35-mm dishes, cultured for 1 day in siRNA medium (DMEM (Gibco) with 10% fetal bovine serum and 2 mM L-Glutamine (Gibco)) to reach 80-90% confluency. Then, cells were transfected with the following siRNAs (Allstars Negative Control siRNA (Cat. No. 1027280), Hs Cbs siRNA mix (Cat. No. SI03058391, SI02777159, SI00001316, SI02777166), Mm Cbs siRNA mix (SI00942606, SI00942620, SI00942613, SI00942599), Hs Cry1 siRNA mix (SI00059430, SI00059416, SI04434780, SI02757370), Mm Cry1 mix (SI02708041, SI02686271, SI02666580, SI02732086) using the Thermo Scientific Lipofectamine RNAiMAX Reagent manufacturer’s instructions. After 2 days, dexamethasone (final 0.2 µM) was added to the medium. After 2 hours of incubation, the medium was replaced with medium (DMEM (Sigma, D2902) supplemented with 10 mM HEPES, 0.35 mg/ml sodium bicarbonate, 100 μg/μl Streptomycin and 100 μg/μl Penicillin (Gibco), 2 mM L-Glutamine (Gibco) and 1 mM luciferin, pH 7.2) and the luminescence was recorded every 10 minutes for 6 days in U2-OS: P(Bmal1)-dluc and NIH 3T3: P(Per1)-dluc cells.

### In vitro Experiments

#### Bimolecular Fluorescence Complementation (Bi-FC)

HEK293T cells in 6-well dishes were transfected with venus plasmids (200 ng CRY1-VC (venus C-terminal, 300 ng CBS-VN (venus N-terminal) and 200 ng pmCherry-C1 plasmids) by using Fugene HD (Promega). At 24 h post-transfection, the cells were visualized using the YFP and mCherry fluorescence microscopy filter sets as described in (Anafi et al., 2014).

#### Co-immunoprecipitation

HEK 293T cells were grown in Dulbecco’s modified Eagle’s medium (DMEM; Invitrogen) supplemented with 10% fetal bovine serum. The day before transfection, 4 × 10^5^ cells were seeded and then a total of 2 μg of equimolar amounts of plasmid DNA constructs (*Cry1*-pcDNA (either wild type, or T1, or R602P) and *Cbs*-pCMV-sport6 (either wild type or I278T) were transfected into the cells with Fugene HD (Promega). Cells were incubated for 48 hours and then DMEM was removed. After washing the cell with PBS 3X, cells were collected by centrifugation at 1000 xg at 4°C for 5 minutes. Pellets were resuspended, dissolved in lysis buffer (50 mM pH 8 Tris-Cl, 150 mM NaCl, 1 mM EDTA and 1% Triton-X supplemented freshly with Protease Inhibitor Cocktail (Sigma)) and sonicated 2 times for 10 seconds with 15% power (Bandelin, Germany). Samples were centrifuged at 14000xg at 4°C for 40 minutes to get rid of cellular debris. Anti-Flag M2-Flag resin or Ni-NTA-His resin equilibration was achieved by washing the resin 2 times with TBS to equilibrate with lysis buffer. Then resin was added to cleared cell lysate and incubated at 4 °C overnight. The next day, beads were washed 3 times with TBS to eliminate non-specific binding, and then beads were boiled with 6x loading dye and elutes were processed for Western blot.

### Generating CRY1 Mutant Library and Lentiviral Cloning Processes

#### Construction of CRY1 Mutant Library Using Degenerate Primers

Random mutagenesis of the last 20 amino acids of the *pACT-mCry1* construct was performed with degenerate reverse primers. The primers used to mutate the odd or even numbered residues are shown in Table S1. PCR was performed in a total volume of 50 μl containing approximately 50 ng of plasmid samples, 20 pmol of each primer, 0.2 mM dNTPs, and 2.5 unit Phusion Taq DNA polymerase. Conditions for the 18 cycles of amplification were 95°C for 30 s, 50°C for 30 s and 68°C for 14 min. Before the first cycle, reaction mixtures were kept at 95°C for 4 min and at the end of the 18th cycle an additional 68°C extension period was applied for 10 min. Samples were then treated with DpnI restriction enzyme to remove the template DNA and transformed into *E coli* DH5α. Transformed cells were plated and selected on appropriate selective media. Mutagenesis results were verified by DNA sequencing carried out by Macrogen (Amsterdam, Netherland).

#### Site Directed Mutagenesis

PCR-based site directed mutagenesis was performed to generate the CRY1-R602P mutant. The PCR reaction contained 30 fmol of DNA, 20 pmol of primers (Supp Table 1), 0.2 mM dNTPs, and 2.5 units of *Pfu* Turbo DNA polymerase. The PCR was carried out for 12 cycles under the following conditions: 40 s at 94 °C, 40 s at 55 °C, and 11 min at 68 °C. The PCR products were digested with DpnI to remove template plasmid DNA and transformed into *E. coli* DH5α. The presence of the site-directed mutations was confirmed by DNA sequencing through the Macrogen sequencing facility.

#### Immunoblotting

Cell lysates were mixed with Western Blot loading dye (final concentrations, 50 mM Tris-Cl, 2 mM EDTA pH 8, 1 % SDS, 1% beta-mercaptoethanol, 8% glycerol, 0.02% bromophenol blue) and boiled for 5 minutes at 95 °C. Samples were subjected to 10% SDS-PAGE to separate proteins. Proteins were transferred to PVDF membrane (Millipore, Immobilon-P-Transfer Membrane) using a BioRad wet-transfer system. After transfer, the membrane was blocked with 5% non-fat milk (Chem Cruz SC2325) prepared in 0.15% TBS-T (Tris-buffered saline with 0.15% tween 20)(Sigma Aldrich Tween 20). After blocking, the membrane was incubated for 45 minutes with either Anti-HIS antibody (Santa Cruz SC-8036), Anti-FLAG antibody (Sigma A9469), Anti-CRY1 antibody (Bethyl), Anti-CBS antibody (Abcam ab54883), Anti-VP16 (Santa Cruz, sc-7545), Anti GAL4 DBD (Santa Cruz, sc-577), Anti ACTIN (Cell Signaling, 8H10D10), Anti α-TUBULIN (Sigma, T9026), or Anti HISTONE-H3 (Abcam, ab1791). Following primary antibody incubation, the membrane was washed 3 times with 0.15% TBS-T for 5 minutes. Then, membranes were incubated with secondary antibodies, either anti-mouse (Santa Cruz SC-358920), or anti-rabbit (abcam ab97051), for 50 min. Next, membranes were washed 3-times with TBS-T for 15 min total. In order to visualize the membrane with chemiluminescent detection, Advansta ECL HRP substrate was used (WesternBright Sirius). Images were taken and quantified by BioRad Chemidoc. Similar procedure was used for gradient blue native-PAGE analysis by preparing the samples without using SDS or any denaturant. Gradient native gel (from 3 to 10 %) was used. The pH of running buffer was adjusted to 7.8 which provides negative charged BSA and CBS proteins, since their isoelectric points are close (5.4 and 5.9 respectively). The full-uncropped images of all Western blots were provided in File S7.

### RNA Isolation and Real-Time PCR

#### RNA Isolation

RNA isolation was performed from both cells and livers with RNeasy Mini Kit (Qiagen Cat. No.74106).

#### cDNA Synthesis

RevertAid First Strand cDNA Synthesis Kit (Thermo) was used for cDNA synthesis.

#### Real-Time PCR

SYBR Green real time PCR mix was used with real time PCR primers of target genes. Real time PCR primers are listed in Table S1.

### Cystathionine-Beta Synthase Enzymatic Activity Detection

#### Liver Lysate Preparation and Determination of CBS Enzymatic Activity

A total 12 mg of mouse liver was homogenized with Dounce homogenizer for 5 minutes in phosphate buffer with pH 6.8 with protease inhibitor. After homogenization, samples were sonicated 6 times for 8 seconds with 90 % power and were centrifuged at 20,0000 xg at 4°C for 45 min. The supernatants, or cell free extracts were used to assay CBS activity. Each reaction consists of 100 mM potassium phosphate buffer (pH 7.4), 10 mM L-cysteine, 2.5 mM pyridoxal-5-phosphate, 30 mM DL-proparglycine (PAG), 100 µg of total protein and 2.5 mM homocysteine in total volume of 800 µl. To minimize the confounding influence of CSE activity on our results, we included DL-proparglycine, an inhibitor of CSE ^28^ in these experiments. A second tube that included 750 µl 1 % zinc-acetate and 50 mg cotton were placed in the reaction tube to capture produced gaseous hydrogen sulfide. After the mixing of the 1,5-2 mg of liver with the reaction mixture, nitrogen gas was applied for 5 minutes inside of the tubes in order to deplete oxygen, and reactions were carried out for 1.5 h at 37 °C. Reaction were terminated by mixing with 500 ul 50% trichloroacetic acid. Then, the pieces of cotton were removed from the outer tubes to measure the amount of H2S. Cotton was transferred to glass tubes containing 3.5 ml distilled water. Then following reagents were added to each sample for color development: 400 µl 20 mM N,N-dimethyl-p-phenylenediamine sulphate prepared in 7.2 M HCl, and 400 µl 30 mM FeCl3 prepared in 1.2 M hydrochloric acid. After color development, the absorbance of each sample at 670 nm was measured in a microplate reader (Synergy H1 Multi-Mode Reader-Biotek) so as to calculate the amount of product.

### Cytosol-Nuclear Fractionation

Wild type animal livers harvested at ZT22 and ZT10 were used for fractionation analysis. 10 mg liver was suspended in 500 ul cytosolic lysis buffer (10 mM HEPES pH 7.9, 10 mM KCl, 0.1 mM EDTA, 0.05 % NP40 with protease inhibitors) and incubated on ice for 10 minutes. The lysate was centrifuged for 3 minutes at 4,000 xg at 4 °C. The resulting supernatant was centrifuged one more time and supernatant collected as the cytosolic fraction. The resulting pellet from lysate centrifugation was suspended in nuclear lysis buffer (20 mM HEPES pH 7.9, 0.4 M NaCl, 1 mM EDTA, 10 % glycerol with protease inhibitors) and sonicated 2 times 10 seconds with 60% power. After centrifugation at 15,000 g for 5 minutes at 4 °C, the supernatant was collected as nuclear fraction. Collected fractions were mixed with Western Blot loading dye. Fractionation efficiency was checked with Western blot by using TUBULIN (Sigma T9026) as cytosolic marker and HISTONE-H3 (abcam ab1791) as nuclear marker.

### Generation of *Cry1^−/−^*, *Cry2^−/−^* and *Cbs ^−/−^* Mice

For the generation and maintenance of *Cry1*^−/−^ or *Cry2*^−/−^ mice, methods were performed as previously described in [17]. Briefly, the original *Cry1*^−/−^ or *Cry2*^−/−^ lines of partial C57BL/6J background were repeatedly backcrossed with C57Bl/6J mice to generate *Cry1*^−/−^ and *Cry2*^−/−^ mice. The homozygous *Cry1*^−/−^ and *Cry2*^−/−^ mice were entrained to a 12 hour light:dark cycle with *ad libitum* food and water. After approximately 8 weeks, mice were sacrificed by asphyxiation with CO2 at either ZT10 or ZT22.

Liver samples were frozen instantly. During the generation of *Cbs*^Zn/Zn^ mice, controlled expression of mutant human CBS rescued neonatal lethality as described by Wang et al [31]. Briefly, *Cbs*^−/+^ heterozygous animals were obtained from Dr. Warren Kruger’s lab). The backcrossed heterozygous animals were provided with 25 mol/L ZnSO4 containing water for induction of the transgene. Then, the siblings were intercrossed and fed with zinc to generate homozygous *Cbs*^Zn/Zn^ mice. The generation and maintenance of the transgenic mice that express the human *Cbs*-I278T gene is described in [31].

### Circadian Behavioral Analysis

Mice were housed with ad libitum food and water in individual cages containing running wheels. Temperature, humidity and light were tightly controlled. 21 wild-type and 26 *Cbs*^Zn/Zn^ mice were entrained for 2 weeks with 12 hours light-dark cycles, and then they were maintained in constant darkness. ClockLab Data Collection (Actimetrics) were used for locomotor activity detection and analysis [19]. The Fourier power spectrum was directly output using ClockLab v 6.0 and used to compute the fraction of total power explained in the circadian period range. The included ClockLab functions for non parametric circadian analysis as described in [32] including rhythm amplitude and inter-daily stability were also used.

### Metabolomics Analysis

Liver tissues (50 mg) were thawed on ice and metabolite extraction was performed with Bligh-Dyer method [67, 68]. Briefly, cell pellets were dissolved in a mixture of 2:1 methanol:chloroform, and then they were vortexed and sonicated. Following the addition of chloroform and water, samples were vortexed and organic and aqueous layers were separated by centrifugation. The aqueous layer was vacuum dried and re-dissolved in an acetonitrile:water (2:1) mixture. After centrifugation to remove fine particulates, the supernatant was transferred to LC-MS certified sample vial for Ultra-performance liquid chromatography.

### Metabolomics/Mass spectroscopy data processing and enrichment analysis

Mass spectroscopy and data analysis was performed as previously described [21]. Briefly, ion counts which were obtained from TargetLynx during mass spectroscopy data processing were processed in version 3.2 R. For the LC column equilibration, a QC (quality control) sample mixture injected at the beginning of every 6 injections to account for instrumental drift. QC data was also used for the normalization of the results. Two samples were excluded based on lab QC standards.

ANOVA analysis was performed in R using the aov() command. The comparison of *Cbs*^Zn/Zn^ and WT involved only two factors: gender and genotype. A three factor ANOVA, including time and the interaction term between time and genotype was used in the analysis of Cryptochrome mutants. Multiple correction was performed using the method of Benjimini and Hochberg as implemented in the p.adjust() command.

Overrepresentation analysis was performed using Metaboanalyst.ca website (v 4.0) [69]. Lists of metabolites with significant ANOVA results (p<0.05 for cryptochrome results, q<.05 for CBS results were using the pathway tool and a custom background comprised of the metabolite identified in the corresponding study. The full, pre-ranked metabolite lists, ordered by -log(p) were separately analyzed using the GSEA Java program [43] and metabolite sets download from the Small Molecule Pathway Database [70] and converted to GMT file format.

### Ethics Statement

The relevant institutional review boards approved all procedures involving the use of experimental animals. Experiments utilizing Cryptochrome mutant mice and corresponding controls were approved by the University of North Carolina Institutional Animal Care and Use Committee. Experiments utilizing CBS transgenic mice were approved the University of Pennsylvania Institutional Animal Care and Use Committee

## Author Contributions

IHK and RA conceptualized and designed the experiments, analyzed the data, and wrote the manuscript. SCK carried out in vitro experiments. EKC carried out CoIP experiments of the truncated CRY1 and CBS. JG carried out circadian studies at cellular level. CPS and AS generated the *Cry1* and *Cry2* knockout animals and wrote the manuscript. HA developed CBS assay. HO carried out mutagenesis studies. SDR, DM and AW carried out metabolomic experiments. WDK generated *Cbs* knockout animals. JBH designed experiments and wrote the manuscript.

## Supplementary Materials

**Fig. S1:**
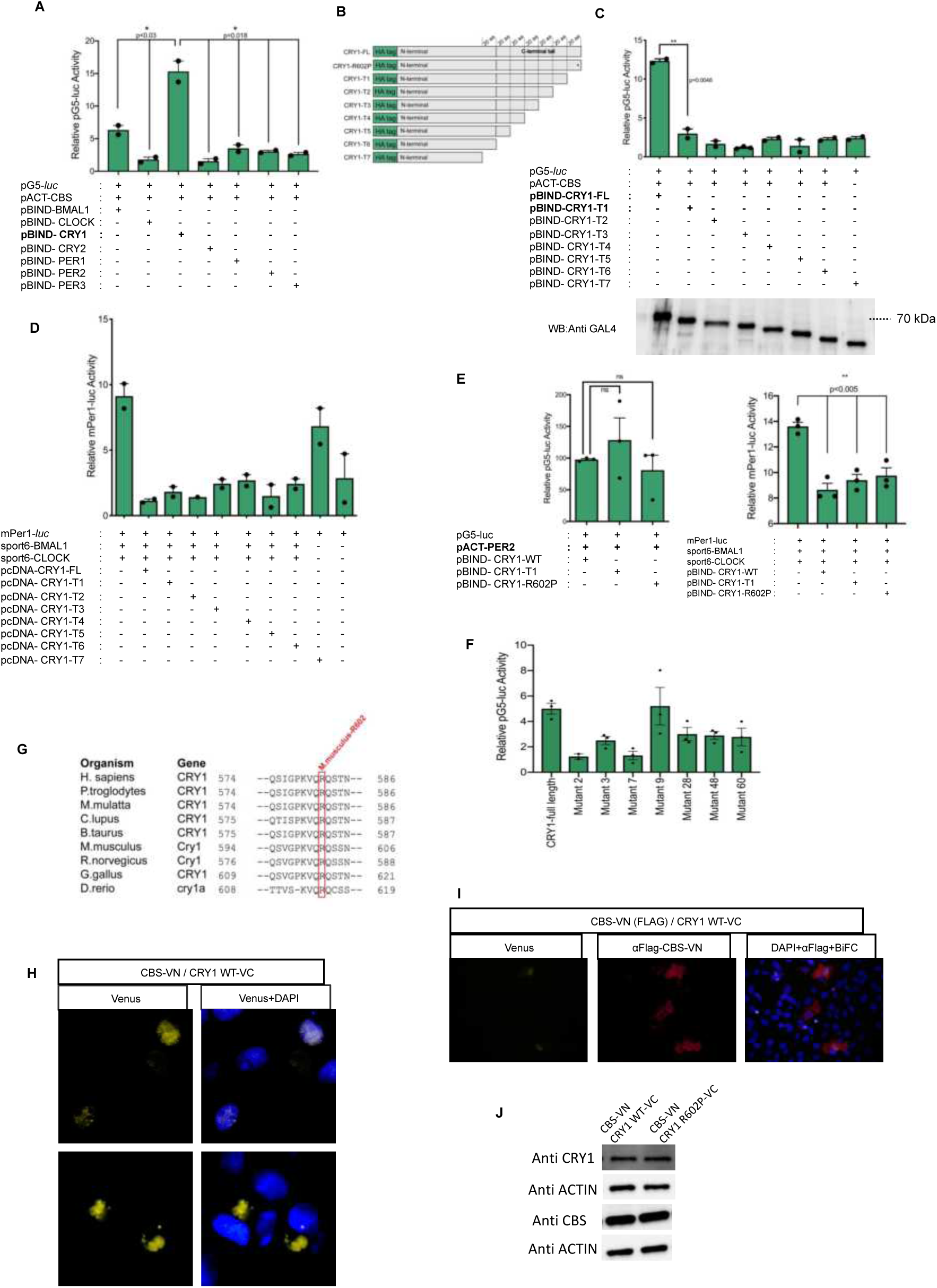
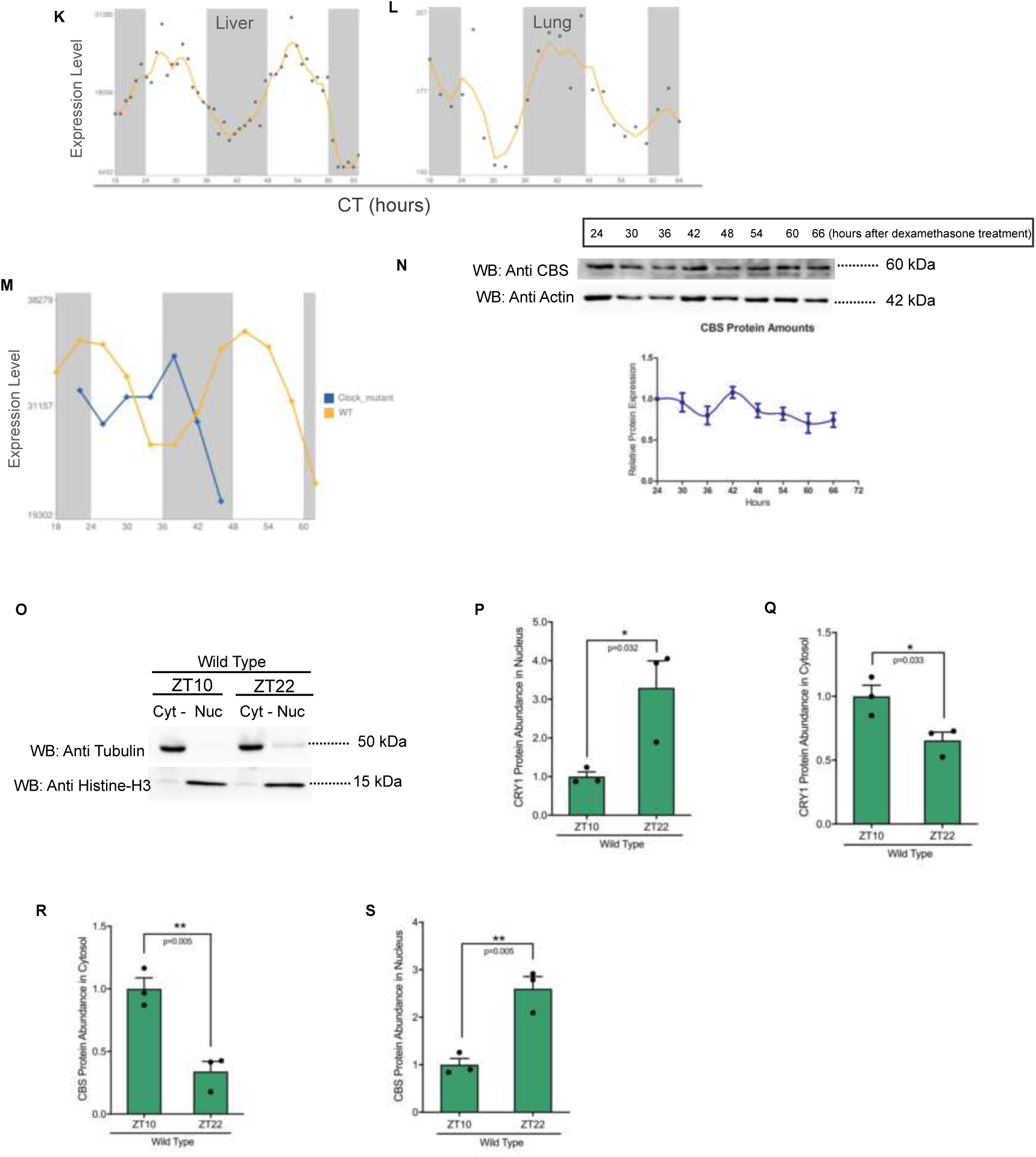
**(A)** Mammalian two hybrid (M2H) interaction analysis of CBS and core clock components. (Bars show the mean luciferase activity ± SEM, n=2, *p<0.03, *p=0.018) **(B)** Schematic representation of CRY1 truncations and mutation. **(C)** M2H analysis of CBS and truncated CRY1 proteins. **Bottom:** Expression of all constructs was confirmed by Western blot using GAL4 antibody (Bars show the mean luciferase activity ± SEM, n=3, **p<0.0048). **(D)** Transactivation activity of BMAL1 and CLOCK of a luciferase reporter from *Per1*-E-Boxes. Full length and truncated CRY1 proteins are able to repress BMAL1-CLOCK transcriptional activity (mean ± SEM, n=2, *p<0.02) (unpaired two-tailed t test). **(E)** The M2H interaction analysis of PER2 and CRY1-WT, CRY1-T1 and CRY1-R602P. Repressive function of wild type, truncated and mutant CRY1 on BMAL1-CLOCK transactivation indicating that all constructs are functional (Bars show the mean luciferase activity ± SEM, n=3). **(F)** M2H interaction analysis of the CBS with wild type CRY1 and mutant CRY1s (Bars show the mean luciferase activity ± SEM, n=3). **(G)** Cross-species alignment of the last 22 amino acids of mouse CRY1. The red box highlights the completely conserved mouse CRY1-R602 in nine diverse species. Multiple alignment analysis was performed on NCBI HomoloGene. **(H)** Bi-molecular Fluorescence Complementation (Bi-FC) assay of CBS and CRY1 showing co-localization at both nucleus and cytoplasm. **(I)** Immunohistochemical confirmation of CBS protein in detected CBS-CRY1 Interaction regions. **(J)** Confirmation of the expressions of CRY1 and CBS in venus plasmids used in Bi-FC studies by Western blot. **(K-L)** CBS mRNA oscillations in liver and lung (*22, 26*). **(M)** Cbs mRNA expression in wild type and clock mutant mice (*28*). **(N)** CBS oscillation shown with Western blot using anti-CBS and anti-Actin antibodies (n=3). **(O)** Fractionation efficiency check with cytosolic (Tubulin) and nuclear (Histone-H3) proteins. **(P-Q)** Quantification of CRY1 protein in nuclear and cytosolic fractions (n=3, mean ± SEM, *p=0.032, * p=0.033). **(R-S)b**Western blot analysis of cytosolic and nuclear CBS levels in wild type ZT10 and ZT22 liver samples. (n=3, mean ± SEM) (unpaired two-tailed t test, **p=0.05).

**Fig. S2:**
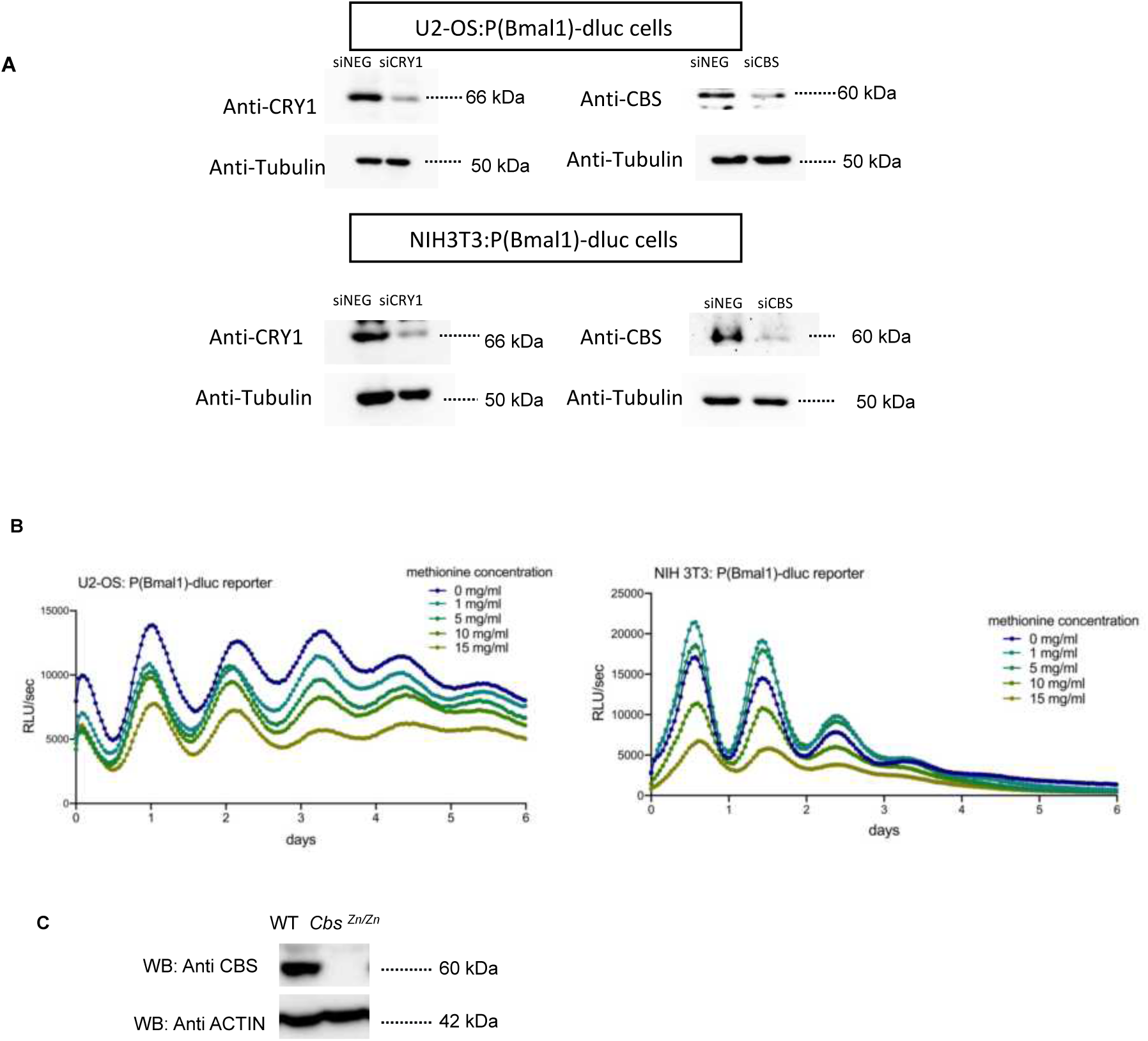
Supplemental data for *in vitro* and *in vivo* circadian experiments. **(A)** Western blot verifying protein level knock down of CBS and CRY1 in U2-OS:(P)Bmal1-dluc and NIH3T3:(P)Bmal1-dluc cells. **(B)** Bioluminescence records of NIH 3T3:P(Bmal1)-dluc and U2-OS:(P)Bmal1-dluc cells under different concentrations of methionine showing no methionine effect on period length. **(C)** The genotypes of the *Cbs*^Zn/Zn^ animals were verified by SDS-PAGE and Western blot with the indicated antibodies using liver samples.

**Fig. S3:**
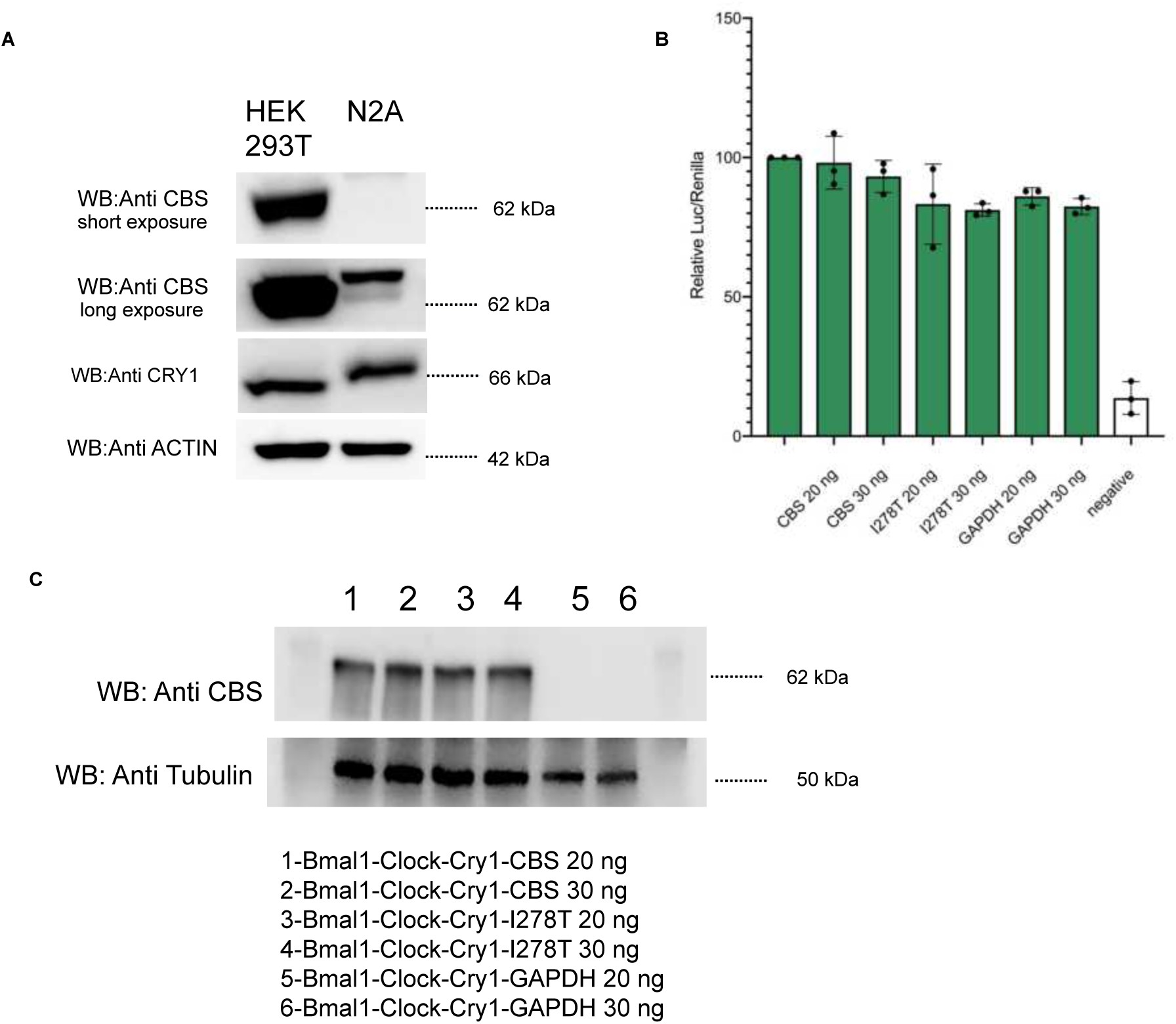
**(A)** Western blot assay showing CRY1 and CBS protein amount in HEK 293T and Neuro-2A (N2A) cells. CBS protein is greatly less in N2A cells. **(B)** Relative Luciferase/Renilla graph showing that CBS WT, CBS-I278T, and GAPDH cannot repress BMAL1-CLOCK transactivation when there is no CRY1. **(C)** Western blot indicating wild type and mutant CBS expressions in repression assay samples from Fig. 3D.

**Fig. S4:**
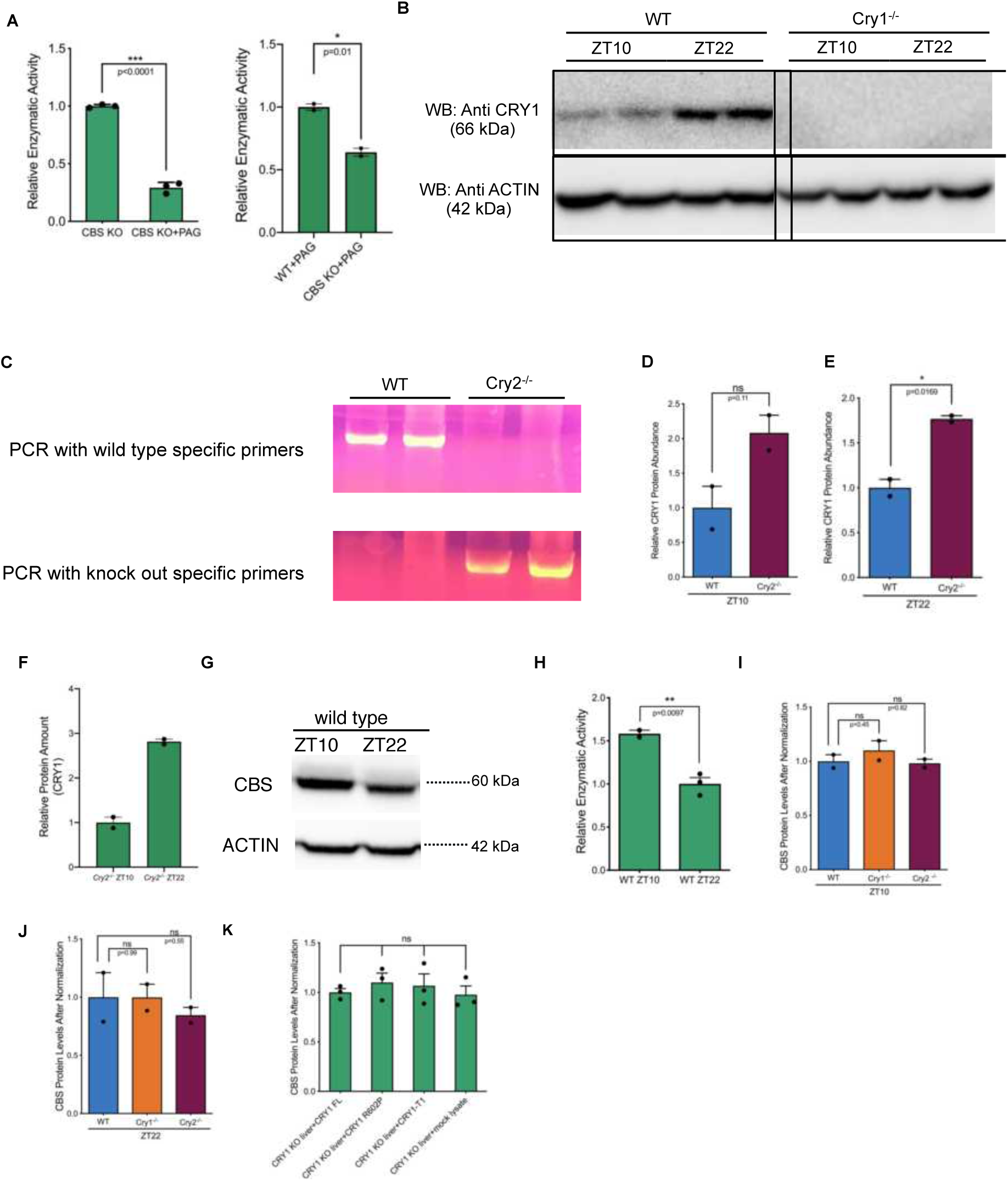
**(A) *Left:***30 mM PAG (DL-proparglycine) inhibits more than 70% CSE (cystathionine gamma lyase) H2S production activity. ***Right:*** CBS in WT animals show significantly more activity than *Cbs*^Zn/Zn^ animals in the presence of CSE inhibitor PAG. (n=2, mean ± SEM, unpaired t-test) **(B)** The genotypes of the *Cry1*^−/−^animals were confirmed with Western blot using liver samples harvested at ZT 10 and ZT 22. CRY1 protein amounts are shown. **(C)** *Cry2*^−/−^genotype confirmation with PCR showing wild type and *Cry2*^−/−^liver genomic DNA samples. Wild type specific and *Cry2*^−/−^ specific primers were used. **(D)** and **(E)** Quantification of CRY1 protein levels in WT and *Cry2*^−/−^liver samples. **(F)** CRY1 protein oscillation in *Cry2*^−/−^liver samples at ZT10 and ZT22. **(G)** CBS protein amounts in wild type ZT 10 and ZT 22 samples. **(H)** CBS enzyme activity in wild type samples at ZT 10 and ZT 22. **(I-J)** Quantification of CBS protein levels in activity assay samples obtained at ZT 10 and ZT 22, (mean ± SEM, ns p=0.45; ns p=0.82 for ZT10) (mean ± SEM, ns p=0.99; ns p=0.55 for ZT22). **(K)** Equal of CBS protein levels in complementation assay samples which was quantified with Western blot.

**Fig. S5:**
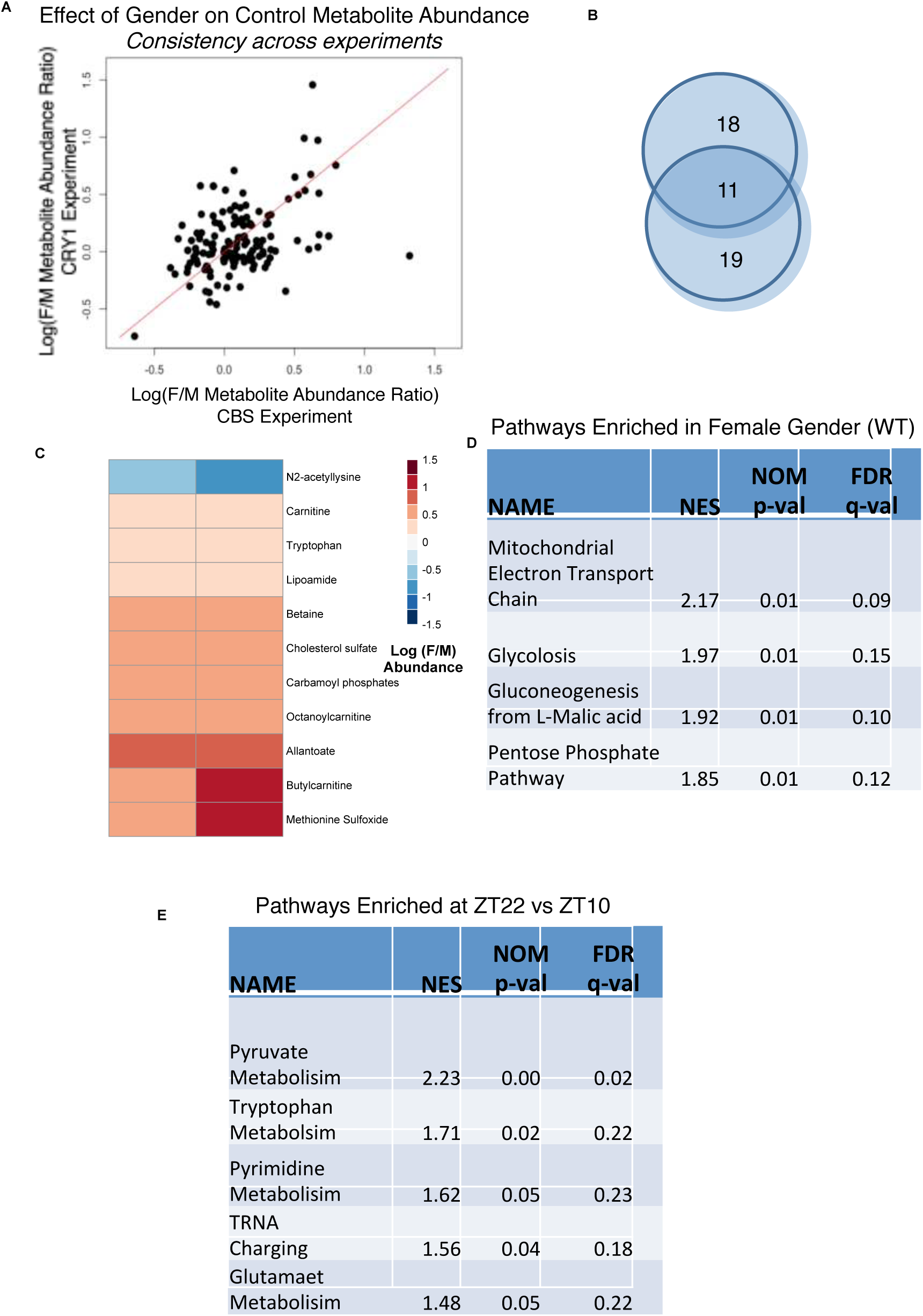
Influence of gender on WT metabolomic results. **(A)** We computed the ratio of mean metabolite abundance in female and male animals using the WT control animals that were used in the two different experiments (*Cbs*^Zn/Zn^ vs WT) and (*Cry1* ^−/−^ vs WT). For each metabolite, the ratio observed in the two experiments is compared. **(B)** ANOVA analyses of the WT control data were used to identify metabolites significantly influenced by gender (FDR<15%) in the two studies. A Venn diagram demonstrates the overlap. **(C)** Heatmap demonstrating the consistency of the gender effect in these overlapping metabolites. **(D)** Metabolites were ranked by the ratio of expression in the female/male mice (using the WT mice from the CRY experiment) set enrichment analysis was used to identify enriched pathways. Pathways with q<.15 are shown. NES is the normalized enrichment score. **(E)** Metabolites were ranked by the ratio of expression in the ZT10 vs ZT22 (using the WT mice of both genders from the CRY experiment). Set enrichment analysis was used to identify enriched pathways for ZT22. No pathways were significantly enriched at ZT10.

**Fig. S6:**
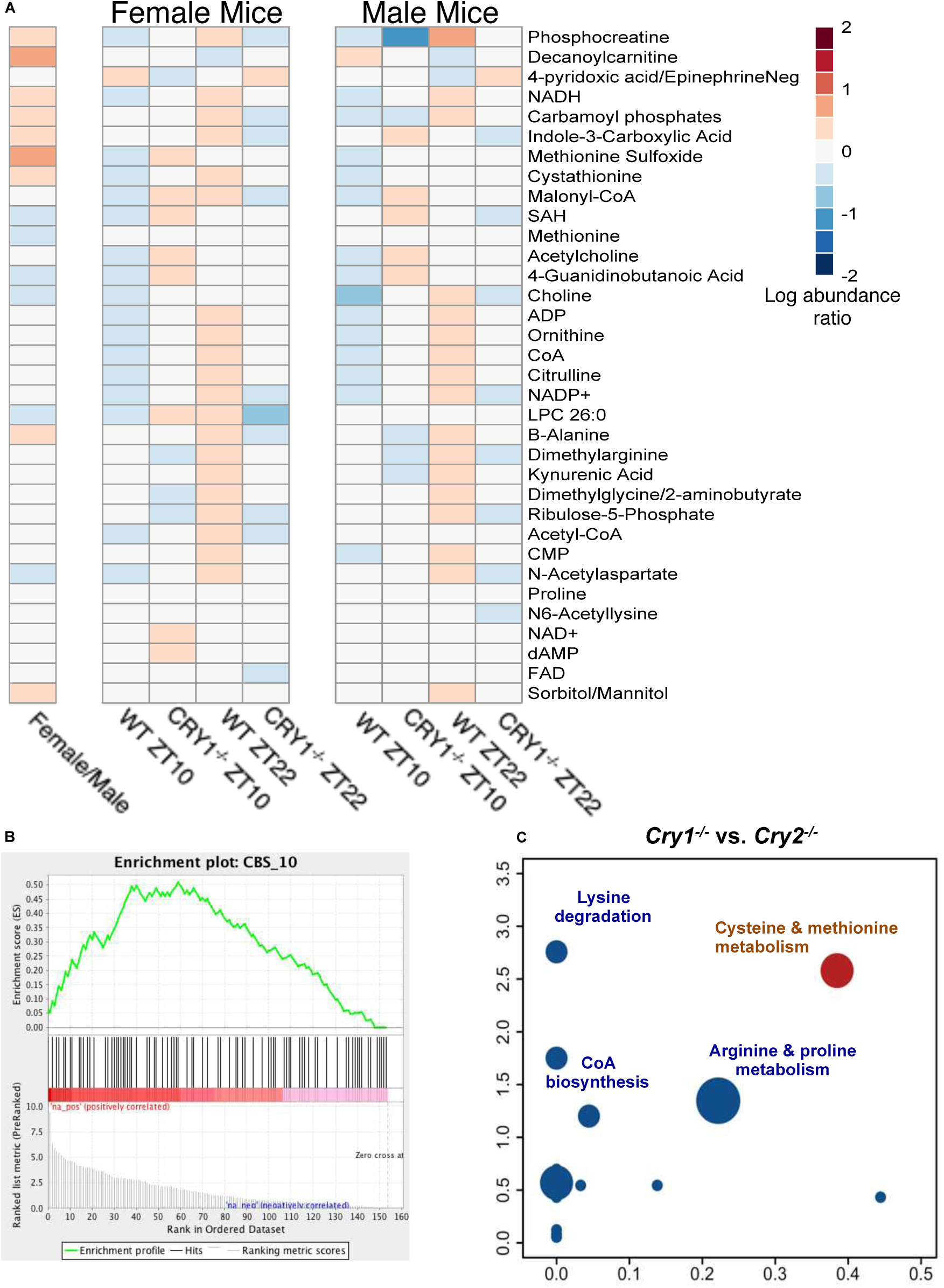
**(A)** Influence of gender on metabolites that are significantly (p<0.05) affected by the *interaction* of cryptochrome genotype and time. These are the same metabolites shown in Fig 6C. The log ratio of mean abundance in female/male mice is shown. Metabolite abundances from samples from male and female mice are shown relative to the mean of samples from that gender (log ratio). The influence of CRY1 genotype and time is consistent across genders. **(B)** Metabolites were ranked by the significance of the interaction term between time (ZT10 vs. ZT22) and genotype (*Cry1^−/−^* versus WT) in the ANOVA. An enrichment plot shows the placement of a custom set of metabolites defined by their responsiveness to CBS deficiency. **(C)** Pathway overrepresentation describing liver metabolites influenced by the interaction of cryptochrome genotype (*Cry1^−/−^* versus *Cry2^−/−^*) and circadian time (p <0.05). Only female *Cry2^−/−^* were available and only female *Cry1^−/−^* mice were used in the analysis.

**Fig. S7:**
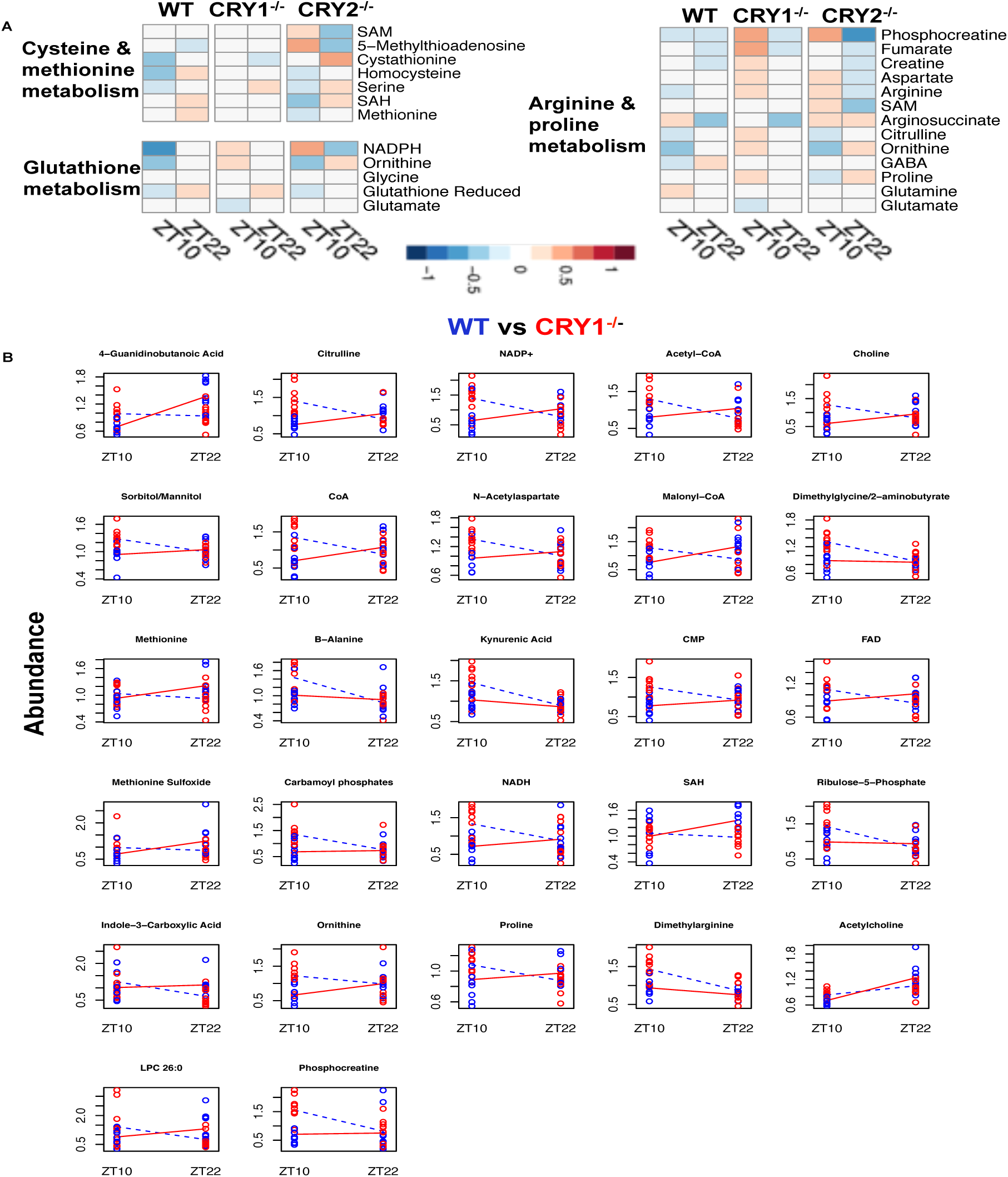
**(A)** Heatmaps showing the influence of time and genotype on the concentration of the listed metabolites. Metabolites are organized by KEGG pathway. **(B)** Interaction plots for individual metabolites significantly affected the interaction of genotype (CRY1 vs WT) and time (ZT10 vs ZT22), (p<0.05). Data from all biological samples are shown. Each biological sample is the average of 3-4 technical replicates. Blue and red circles denote data from WT and *Cry1^−/−^* mice respectively. Lines connect mean values from ZT10 to ZT22.

**Table S1:**
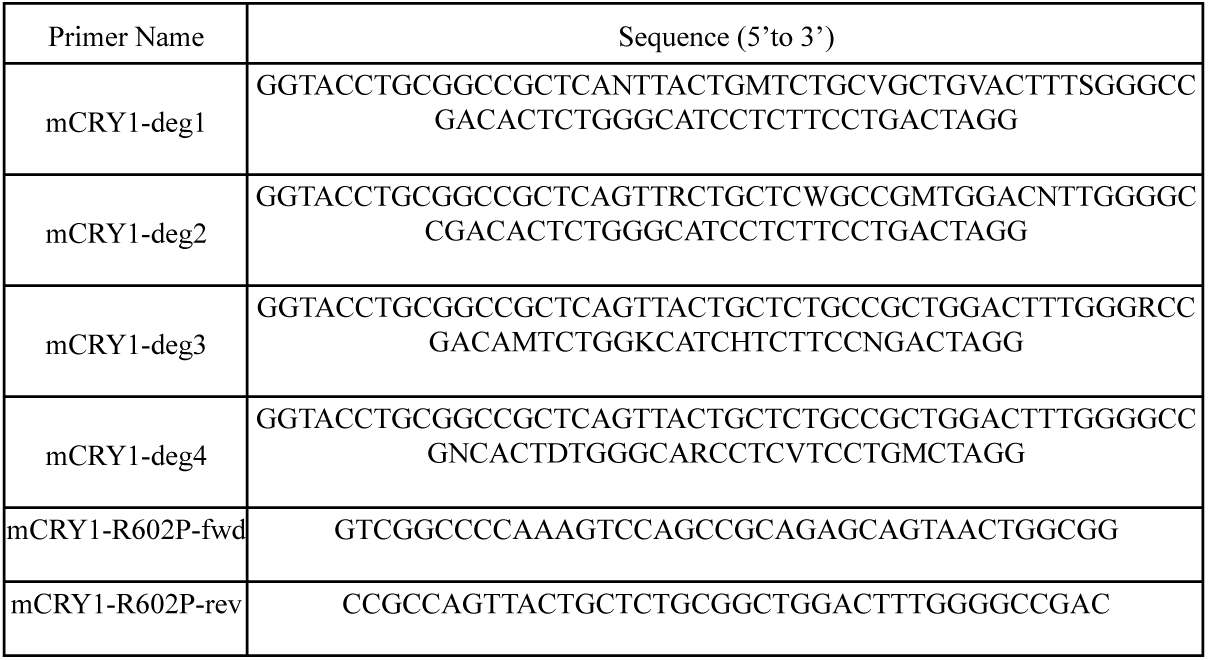
Degenerate and site directed PCR primers used to mutate the CRY1 C-terminal region.

**Table S2:**
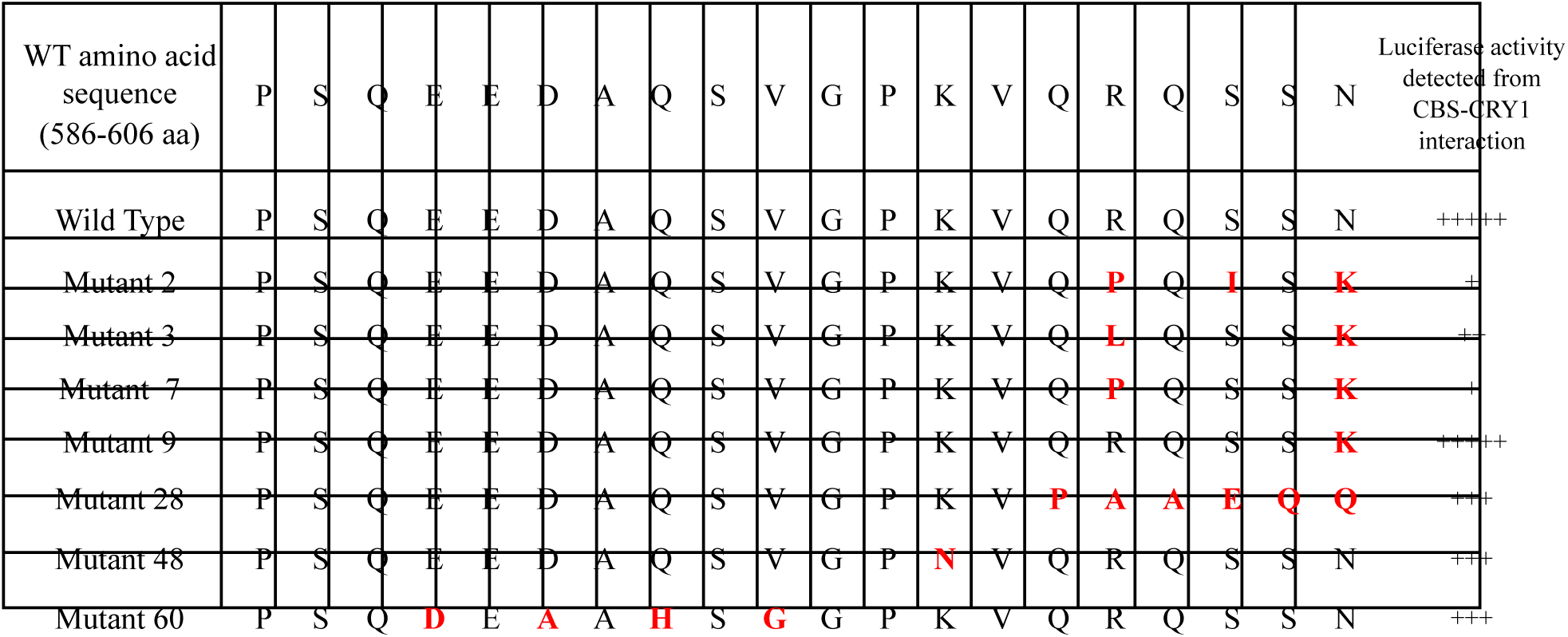
Last 20 amino acids are shown for some of the mutated CRY1 constructs. Mutations are indicated with red. The last column shows the luciferase activity yielded by CBS and CRY1 interaction. Each “+” represents fold of interaction detected by mammalian two hybrid assay (Fig. *S1F*). A common theme for the degenerate mutants exhibiting a significant decrease in luminescence signal intensity was the presence of a mutation in R602. R602P mutations disrupted and in some cases abolished CRY1-CBS interaction.

**Table S3:**
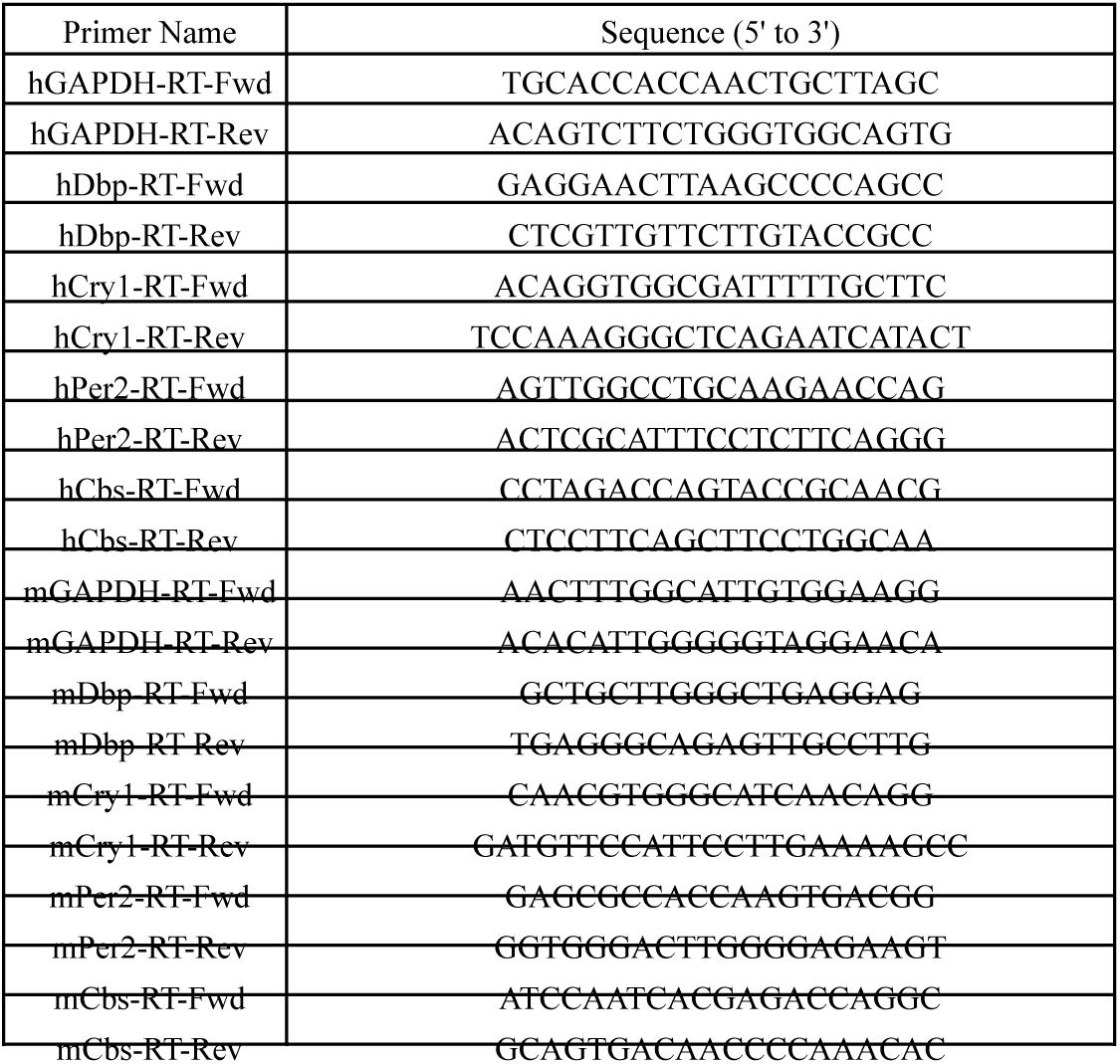
Real-time PCR primer list for both human and mouse *Dbp*, *Cry1*, *Per2* and *Cbs*.

